# Sex Differences in Brain Tumor Glutamine Metabolism Reveal Sex-Specific Vulnerabilities to Treatment

**DOI:** 10.1101/2021.09.29.461531

**Authors:** Jasmin Sponagel, Jill K. Jones, Cheryl Frankfater, Shanshan Zhang, Olivia Tung, Kevin Cho, Kelsey L. Tinkum, Hannah Gass, Elena Nunez, Douglas R. Spitz, Prakash Chinnaiyan, Jacob Schaefer, Gary J. Patti, Maya S. Graham, Audrey Mauguen, Milan Grkovski, Mark P. Dunphy, Simone Krebs, Jingqin Luo, Joshua B. Rubin, Joseph E. Ippolito

## Abstract

Sex differences in normal metabolism are well described, but whether they persist in cancerous tissue is unknown. We assessed metabolite abundance in glioblastoma surgical specimens and found that male glioblastomas are enriched for amino acids, including glutamine. Using PET imaging, we found that gliomas in male patients exhibit significantly higher glutamine uptake. These sex differences were well-modeled in murine transformed astrocytes, in which male cells imported and metabolized more glutamine and were more sensitive to glutaminase 1 (GLS1) inhibition. The sensitivity to GLS1 inhibition in males was driven by their dependence on glutamine-derived glutamate for α-ketoglutarate synthesis and TCA cycle replenishment. Females were resistant to GLS1 inhibition through greater pyruvate carboxylase-mediated TCA cycle replenishment. Thus, clinically important sex differences exist in targetable elements of metabolism. Recognition of sex-biased metabolism is an opportunity to improve treatments for all patients through further laboratory and clinical research.

## Introduction

Cancerous growth requires extensive adaptations at the cellular, tissue, and systemic levels. Some adaptations create not only a growth advantage but also vulnerabilities that can be targeted. Targetable cancer cell adaptations include dependency on certain nutrients (e.g., glucose and glutamine), ATP production, reactive oxygen species (ROS) management, and macromolecule synthesis. These adaptations are collectively referred to as metabolic reprogramming (Pavlova & Thompson, 2016).

Metabolic reprogramming affects carbohydrates, amino acids, fatty acids, and mitochondrial function. Importantly, each of these elements of metabolism exhibit sex differences at the organismal, tissue, and cellular level from the moment of fertilization throughout adulthood and across many species (Rubin et al., 2020). Furthermore, sex-specific metabolic phenotypes have been linked to sex differences in diabetes and cardiovascular disease (Murphy et al., 2017; Tramunt et al., 2020) as well as cancer. In renal cell carcinoma, colon cancer, and low-grade glioma, certain metabolic phenotypes correlate with poor survival in one sex but not the other (Cai et al., 2020; Ippolito et al., 2017; Lopes-Ramos et al., 2020; Nguyen et al., 2018), suggesting that metabolic reprogramming and its effects on survival can differ in male and female cancers. This may be especially important as sex differences in cancer incidence and mortality rate are evident throughout the world, across all ages, and in many different cancer types (Dong et al., 2020; Edgren et al., 2012; Siegel et al., 2021; Zheng et al., 2019). To date, the molecular mechanisms underlying sex-specific metabolic phenotypes in cancer and how these differences might affect metabolic treatment approaches in male and female patients remain unknown.

Here, we show that male and female glioblastoma (GBM) surgical specimens differ in their metabolic profile and that glutamine uptake is greater in male patient glioma and male murine transformed astrocytes. Furthermore, inhibition of glutaminase 1 (GLS1) is more effective against male transformed astrocytes and GLS1 inhibition affects the TCA cycle and glutathione in a sex-specific manner. Moreover, while male transformed astrocytes are more dependent on glutathione to maintain their redox balance, the underlying reason for their sensitivity to GLS1 inhibition is decreased TCA cycle influx from glutamine-derived glutamate. Lastly, we show that resistance to GLS1 inhibition in female cells results from pyruvate carboxylase (PC)-mediated carbon flux from glucose into the TCA cycle. Understanding how metabolism differs in male and female GBM will not only reveal novel elements of GBM biology but will also help stratify patients for metabolic treatment approaches and improve survival for both men and women with GBM and other cancers.

## Results

### Sex differences in GBM metabolite abundance parallel those found in serum

Physiologic sex differences in metabolism are well described. A large-scale metabolomics analysis of over 500 metabolites in blood serum of 1756 participants found that 1/3 of all metabolites showed significant sex differences (Krumsiek et al., 2015). Furthermore, amino acid and carbohydrate pathways were most significantly enriched in males. To determine whether such sex differences persist in cancerous tissue, we assessed metabolite abundance in a previously published dataset of 757 metabolites in 44 male and 32 female newly diagnosed GBM surgical specimens (Prabhu et al., 2019).

We removed metabolites that were constant across all samples, log transformed individual metabolite data, and derived a z-score for each metabolite. In a pathway-based approach, we averaged the z-scores across metabolites that belonged to the same pathway to represent metabolic pathway activity. First, we assessed the “super-pathways,” including lipids, carbohydrates, amino acids, xenobiotics, nucleotides, energy, peptides, and cofactors and vitamins. While none of these were significantly enriched in males versus females at the 5% false discovery rate (FDR) level, we found that all super-pathways were male-biased (Fig. 1A). Similar to the published serum metabolite data, the amino acid and carbohydrate super-pathways were most strongly enriched in males (FDR adjusted p-value q=0.11). Each super-pathway was then subdivided into two or more “sub-pathways.” While no amino acid or carbohydrate sub-pathways were significantly enriched in males versus females at the 5% FDR level, we found that similar to the super-pathways, most sub-pathways were enriched in male GBM (Fig. 1B). Notably, the lipids super-pathway was the only pathway that exhibited a female-bias in multiple sub-pathways, again paralleling the findings in human serum. Although underpowered to detect significant sex differences in pathway enrichment, these data suggest that sex differences in metabolite abundance in GBM tissue and healthy serum are similar.

**Figure 1:**
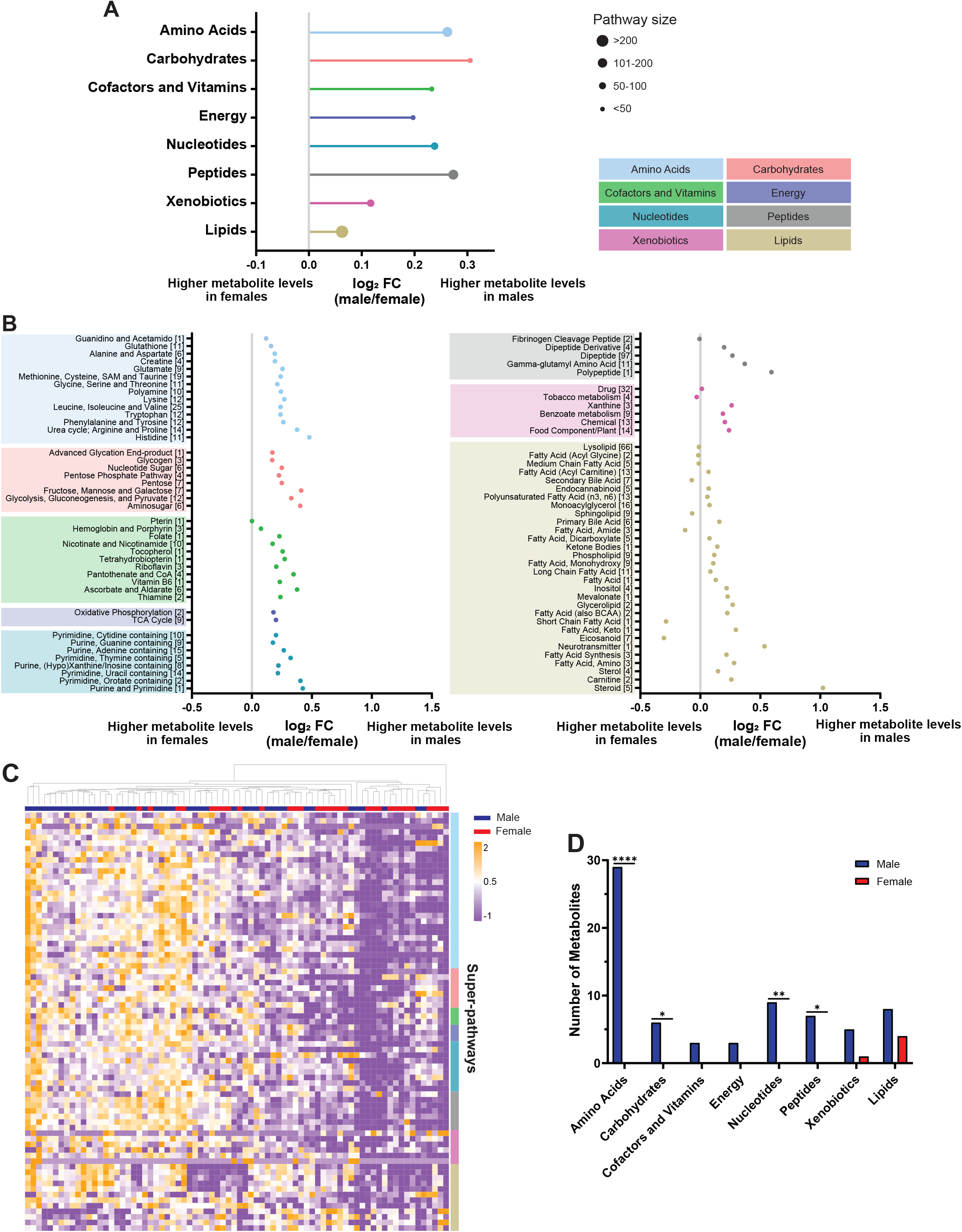
Sex differences in GBM metabolite abundance parallel those found in serum. **A,** Log_2_ fold change (log_2_ FC, indicating male/female mean difference) of metabolic super-pathways in male (n=44) and female (n=32) GBM surgical specimens. Log_2_ FC>0 and log_2_ FC<0 corresponds to super-pathway enrichment in male and female GBM, respectively. Pathways are organized based on Storey’s FDR adjusted q-values, with the smallest q-value at the top. The size of the dots represents the number of metabolites in each super-pathway. **B,** Log_2_ FC of metabolic sub-pathways in GBM surgical specimens. Log_2_ FC>0 and log_2_ FC<0 corresponds to sub-pathway enrichment in male and female GBM, respectively. Sub-pathways are organized based on Storey’s FDR adjusted q-values, with the smallest q-value at the bottom of each super-pathway. The numbers in brackets refer to the number of metabolites in each sub-pathway. **C,** Hierarchical clustering of the top 75 metabolites (determined by greatest mean difference) of male and female GBM surgical specimens. Patient specimens and metabolites are represented in columns and rows, respectively. Rows were sorted based on super-pathway classification (indicated by color on the right). **D,** Distribution of the top 75 metabolites with differential abundance between male and female GBM surgical specimens into super-pathways. *p<0.05, **p<0.01, ****p<0.0001 (two-tailed Fisher’s exact test).

Next, we assessed sex differences in individual metabolite abundance. We selected 75 metabolites with the highest mean difference between male and female GBM surgical specimens and performed unsupervised hierarchical clustering (Fig. 1C). Most patient specimens with high metabolite abundance were males. To further evaluate this observation, we performed unbiased k-means clustering (n=2) (Ippolito et al., 2017) on the same data (Fig. S1A). Cluster 1 included the low, and cluster 2 the high metabolite abundance groups. We found that females and males were associated with clusters 1 and 2, respectively (p=0.0018; Fisher’s exact test) (Fig. S1B), indicating that differences in metabolite abundance allow for sex-specific clustering of GBM. Lastly, we assigned each of the 75 metabolites to a super-pathway and assessed whether metabolites of certain super-pathways were enriched in males or females. Most metabolites belonged to the amino acid super-pathway and were significantly enriched in males (p<0.0001, Fig. 1D). Similar to the sub-pathway analysis, most of the metabolites enriched in females belonged to the lipids super-pathway. Together, these data indicate that male and female GBM differ in their metabolic profile. Super-pathway and individual metabolite analyses identified amino acids as key metabolites enriched in males.

### Males exhibit greater glutamine uptake in glioma and brain tissue

The clinical metabolite data identified amino acids as a super-pathway enriched in male GBM. Interestingly, the metabolite with the greatest mean difference (i.e., male-biased) was pyroglutamine, a cyclic derivative of glutamine. Glutamine was also enriched in males, suggesting that glutamine metabolism might differ between male and female gliomas. This has clinical importance, as inhibition of glutamine metabolism is being evaluated in cancer clinical trials (NCT02071927, NCT04250545, NCT03528642, NCT03965845). To further investigate sex differences in glioma glutamine utilization, we analyzed published and unpublished [^18^F]fluoroglutamine ([^18^F]FGln) positron emission tomography (PET) data (Dunphy et al., 2018; Grkovski et al., 2020; Venneti et al., 2015) from a heterogeneous group of male and female glioma patients separately (Table S1). PET imaging showed marked [^18^F]FGln uptake that was associated with the known brain tumor lesion on magnetic resonance imaging (MRI) in both male and female patients (Fig. 2A). To quantify sex differences in [^18^F]FGln uptake, we measured maximum standardized uptake value normalized to lean body mass (SUL_max_), a method for normalizing tracer uptake in tissues to sex-specific differences in body composition (Tahari et al., 2014). Concordant with our metabolomic profiling, male gliomas exhibited significantly higher [^18^F]FGln uptake than female gliomas (Fig. 2B).

**Figure 2:**
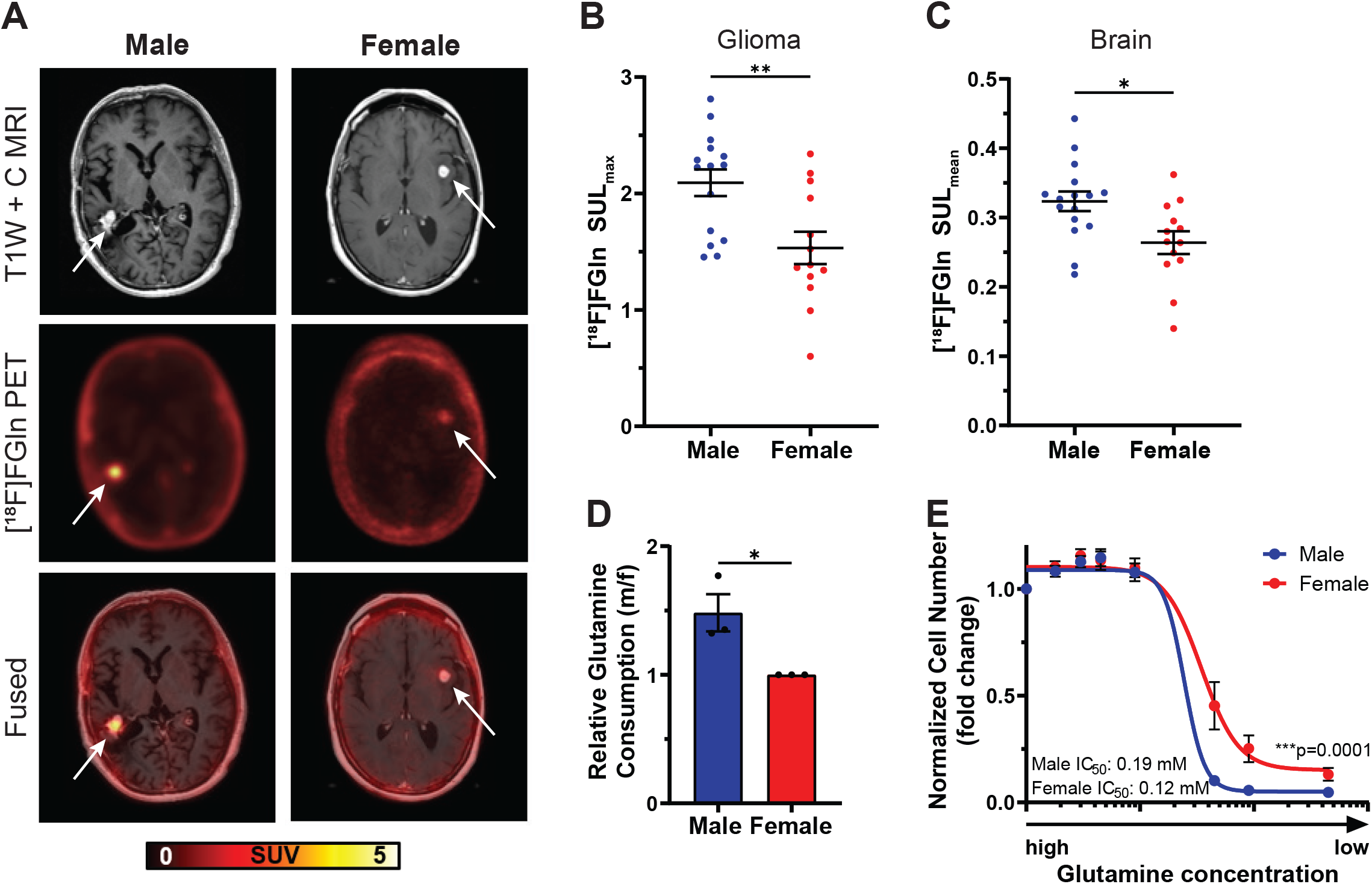
Males exhibit greater glutamine uptake in glioma and brain tissue. **A,** Representative T1W contrast MRI image, [^18^F]fluoroglutamine ([^18^F]FGln) PET image, and fused image of a male and female high-grade IDH wild-type glioma patient. Representative images were chosen based on tumor location, MRI contrast enhancement, and tumor size. Tumor location is indicated by a white arrow. **B-C,** Quantification of [^18^F]FGln uptake in human gliomas (**B**) and normal grey matter (**C**). **D**, Relative glutamine consumption in transformed astrocytes. **E,** Glutamine dose-response curves of transformed astrocytes. Data are mean +/- SEM (B-C) of n=15 male and n=13 female patients or mean +/- SEM of n=3/sex (three independent experiments, one male and female cell line) (D); *p<0.05, **p<0.01 (t-test). Data are mean +/- SEM of n=9/sex (three independent experiments, three separate male and female cell lines); ***p=0.0001 two-way ANOVA (E).

Sex differences in glutamine uptake persisted even when the analysis was restricted to IDH wild-type (WT) glioma patients (Fig. S2A), indicating that the difference in glutamine uptake was not skewed by unequal numbers of IDH mutant or WT gliomas in the male or female patient groups. A similar trend was evident in the IDH mutant patients, but the sample size was too small for a statistically reliable conclusion (Fig S2B). When we restricted the analysis to either GBM (Fig. S2C) or all high-grade gliomas (HGG) (Fig. S2D), we continued to see a similar trend. While in GBM patients, the sample size was too small for a statistically reliable conclusion, HGG showed a strong trend (p=0.06) towards higher uptake in males, indicating that these sex differences were not entirely driven by unequal distribution of HGG and low-grade gliomas (LGG) in the male or female patient groups. Due to small sample size, we could not analyze the LGG patient cohort by itself.

To determine whether sex differences in glutamine uptake were restricted to tumor tissue, we measured [^18^F]FGln uptake in the grey matter of the contralateral cortex of the same patients. The mean SUL (SUL_mean_) value of [^18^F]Gln was significantly higher in male versus female brains (Fig. 2C). Together, these data suggest that sex differences in physiologic glutamine metabolism may be recapitulated in glioma.

To explore the mechanisms underlying sex differences in glutamine metabolism, we took advantage of a previously developed *in vitro* transformed mouse astrocyte model involving combined loss of Nf1 and p53 (Kfoury et al., 2018; Sun et al., 2014). We assessed glutamine consumption in male and female transformed astrocytes and found that male cells consumed ≈1.5-fold more glutamine than female cells (Fig. 2D). Increased glutamine consumption suggests that male cells are more dependent on glutamine for growth. To test this hypothesis, we cultured male and female cells under varying glutamine concentrations and measured cell number. Lowered glutamine levels led to greater reductions in cell number in males compared to females (Fig. 2E). These data indicate that sex differences in glutamine metabolism are present across species and that our transformed astrocyte model is appropriate to study the molecular mechanisms underlying sex differences in glutamine dependency in glioma.

### Male transformed astrocytes exhibit higher glutamine utilization

To delineate specific glutamine requirements in male and female transformed astrocytes, we performed [^13^C_5_ ^5^N_2_]glutamine ([C_5_ N_2_]Gln) tracing experiments. Upon cellular uptake, glutamine is rapidly metabolized to glutamate by the enzyme glutaminase (GLS). The nitrogen that is released during this reaction is required for nucleotide synthesis (Fig. 3A). We assessed nitrogen label incorporation from [^13^C_5_ ^5^N_2_]Gln into the purine guanine and the pyrimidine thymine using solid-state NMR and found that male cells incorporated approximately 20% more nitrogen from [^13^C_5_ N_2_]Gln into nucleotides than female cells (Fig. 3B).

**Figure 3:**
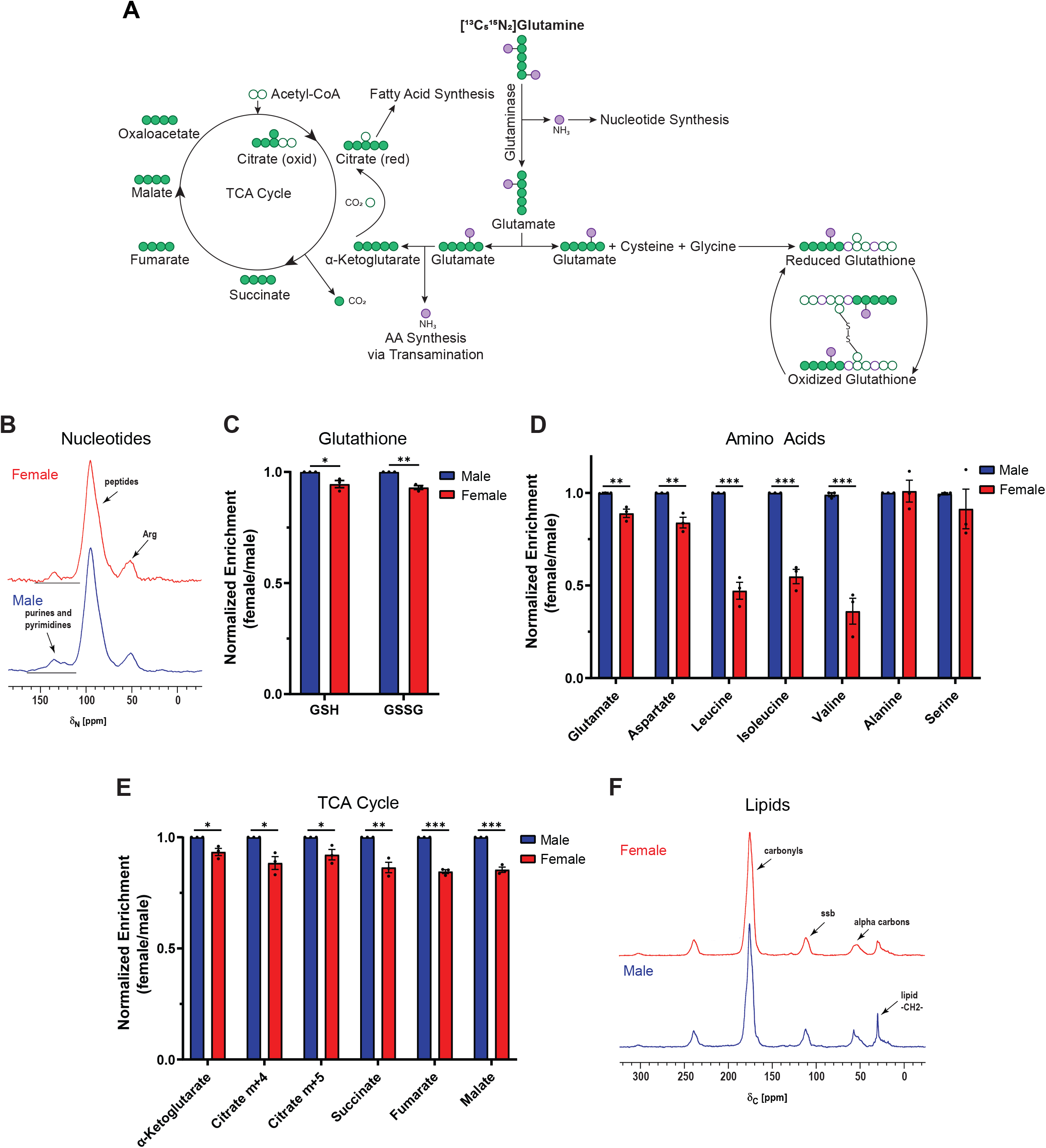
Male transformed astrocytes exhibit higher glutamine utilization. **A,** Schematic of major downstream metabolic pathways of glutamine. **B-F**, Label incorporation of [^13^C_5_ N_2_]glutamine ([^13^C_5_ N_2_]Gln) in transformed astrocytes. Label incorporation into nucleotides (**B**) and lipids (**F**) was measured via solid-state NMR. 50.3-MHz ^15^N NMR spectra (**B**) and 125.7-MHz cross-polarization magic-angle spinning ^13^C NMR spectra (**F**) are shown. Label incorporation into reduced (GSH) and oxidized (GSSG) glutathione (**C**) was measured via LC/MS. Prominent (fully labeled) isotopologues for GSH (m+5) and GSSG (m+10) are shown. Label incorporation into amino acids (**D**) and TCA cycle intermediates (**E**) was measured via GC/MS. Prominent isotopologues for glutamate (m+6) and all other amino acids (m+1) are shown. Prominent labels (one turn of the TCA cycle) for α-KG (m+5), citrate (oxidative TCA cycle (m+4); reductive TCA cycle (m+5)), succinate (m+4), fumarate (m+4), and malate (m+4) are shown. Data are mean +/- SEM of n=3/sex (three independent experiments, one male and female cell line). *p<0.05, **p<0.01; t-test (C). *q<0.05, **q<0.01, ***q<0.001; t-test, FDR adjusted p-values (D-E).

As a key component of glutamine metabolism, glutamate feeds into two major downstream pathways, glutathione and α-ketoglutarate (α-KG) synthesis. Together with glycine and cysteine, glutamate can form glutathione. Glutathione is the most abundant small molecular weight thiol in cells and acts as a free radical scavenger as well as a substrate for electrophile detoxification. Thus, glutathione plays a critical role in maintaining cellular redox balance and has been associated with chemotherapy resistance in cancer (Bansal & Simon, 2018). Glutathione exists primarily in its reduced form (GSH) in cells, but under oxidizing conditions it is converted to glutathione disulfide (GSSG). We assessed carbon label incorporation from [^13^C_5_ N_2_]Gln into GSH and GSSG using LC/MS and found that male cells incorporated significantly more carbons from [^13^C_5_ N_2_]Gln into both pools than female cells (Fig. 3C).

Glutamate can also be converted into α-KG via oxidative deamination. The liberated amino group from this reaction serves as the donor for amino acid synthesis via transamination. We assessed nitrogen label incorporation from [^13^C_5_ N_2_]Gln into multiple amino acids using GC/MS. Aspartate and particularly the branched-chain amino acids (BCAAs) leucine, isoleucine, and valine showed higher nitrogen label incorporation from [^13^C_5_ N_2_]Gln in male cells compared to female cells (Fig. 3D). This was not the case for serine and alanine, highlighting a potentially unique nitrogen requirement from glutamine for BCAA synthesis in male cells.

In addition to providing an amino group, the conversion from glutamate to α-KG replenishes the TCA cycle. We assessed incorporation of [^13^C_5_ N_2_]Gln into TCA cycle metabolites using GC/MS and found that male cells incorporated significantly more carbons from glutamine into the TCA cycle than female cells (Fig. 3E). Citrate is an important substrate in fatty acid synthesis. Glutamine can replenish the cellular citrate pool either via the oxidative direction (m+4), or the reductive direction (m+5) of the TCA cycle (Fendt et al., 2013). Both citrate isotopologues had significantly higher label incorporation in male cells, suggesting that males utilize more glutamine in both directions of the TCA cycle. This indicates that males might utilize more glutamine for fatty acid synthesis. We assessed [^13^C_5_ N_2_]Gln incorporation into lipids using solid-state NMR and found that in male cells carbon incorporation from glutamine is approximately 2-fold higher than in female cells (Fig. 3F). This extensive accounting of glutamine utilization in male and female transformed astrocytes shows that males are more dependent on glutamine for downstream cellular processes than their female counterparts.

### GLS1 expression and dependency is greater in male transformed astrocytes and gliomas

Targeting glutamine metabolism continues to evolve as an attractive cancer treatment. Targeting GLS, which hydrolyzes glutamine to glutamate, was effective in pre-clinical cancer models (Boysen et al., 2019; Jin et al., 2020; Momcilovic et al., 2018; Oizel et al., 2020; Wicker et al., 2021) and its clinical efficacy is currently being evaluated. The GLS family consists of two members, GLS1 and GLS2, which are expressed in the mitochondria and the cytosol, respectively. In brain tissue, GLS1 is expressed at higher levels than GLS2 (Cardona et al., 2015). Accordingly, the predominant enzyme in gliomas is GLS1 (Szeliga et al., 2005). To determine if there was a sex difference in GLS expression, we examined mRNA expression levels in male and female GBM samples from the TCGA Firehose Legacy GBM patient dataset. We found that in both, male and female IDH WT GBM, GLS1 expression levels were significantly higher than GLS2 expression levels (Fig. S3A). Interestingly, GLS1, but not GLS2, was expressed at significantly higher levels in male versus female GBM (Fig. 4A and S3A). Next, we assessed GLS mRNA expression levels in male and female human glioma cell lines from the Broad Institute Cancer Cell Line Encyclopedia and found that GLS1 expression was significantly higher in male glioma cell lines (Fig. 4B). GLS2 expression was low and did not differ by sex (Fig. S3B). These data suggest that GLS1 may play an important role in the sex differences we observed in glutamine metabolism in glioma.

**Figure 4:**
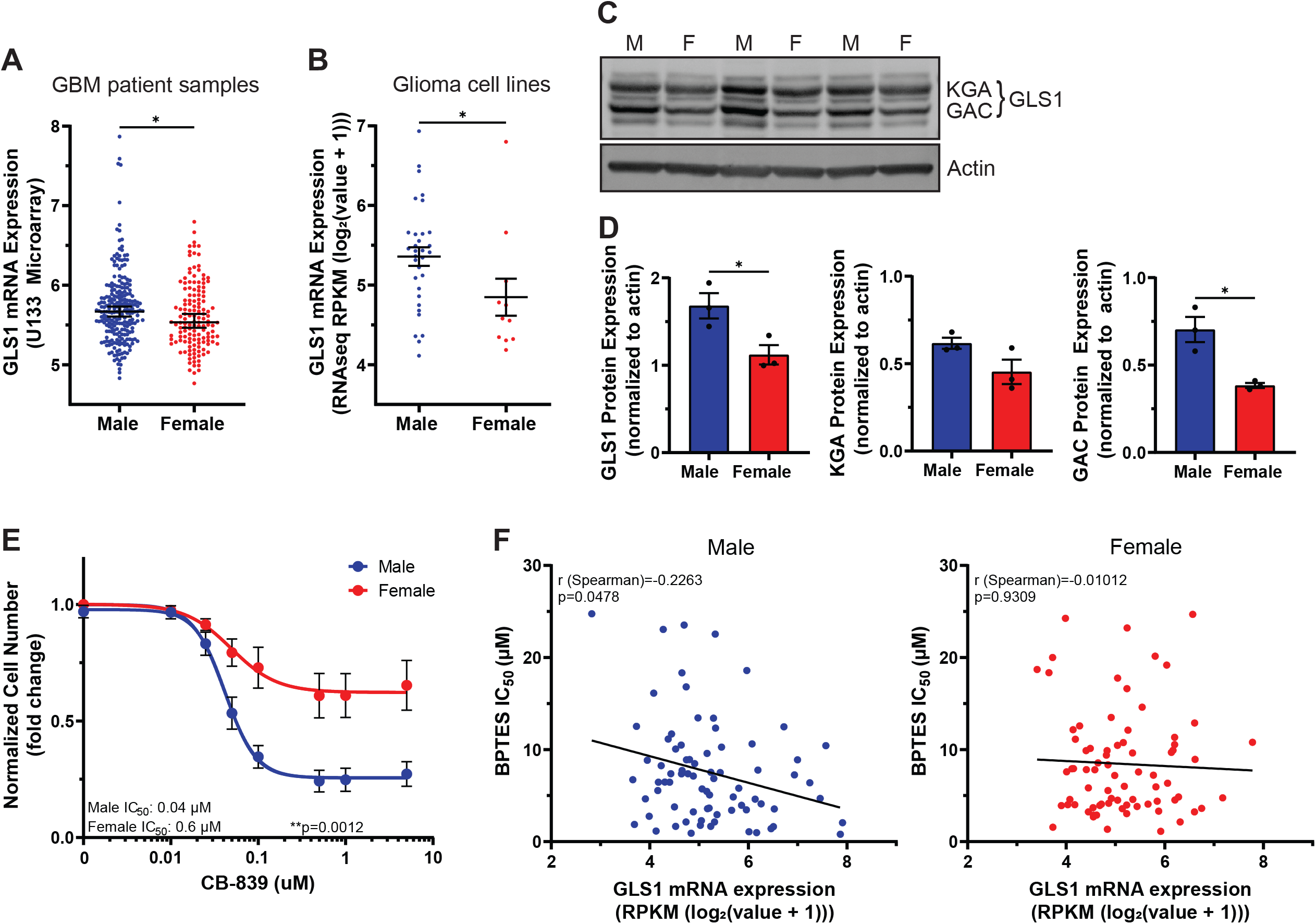
GLS1 expression and dependency is greater in male transformed astrocytes and gliomas. **A-B**, GLS1 mRNA expression levels in n=232 male and n=139 female IDH wild-type GBM patient samples (**A**) and n=32 male and n=11 female human glioma cell lines (**B**). **C,** Western Blot showing the two GLS1 isoforms (KGA and GAC) in transformed astrocytes from three independent experiments. **D**, Western blot quantification of GLS1 and its isoforms KGA and GAC. Expression values are normalized to β-actin. **E**, CB-839 dose-response curves of transformed astrocytes. **F,** Nonparametric Spearman correlation between GLS1 mRNA expression levels and BPTES IC_50_ values in n=77 male (left) and n=76 female (right) tumor cell lines. Data are median +/- 95% CI; *p<0.05; Mann-Whitney test (A-B). Data are mean +/- SEM of n=3/sex (three independent experiments using one male and female cell line); *p<0.05; t-test (D). Data are mean +/- SEM of n=9/sex (three independent experiments, three separate male and female cell lines); **p=0.0012; two-way ANOVA (E).

GLS1 protein expression was also significantly higher in male versus female transformed astrocytes (Fig. 4C-D). GLS1 exists in two isoforms, kidney glutaminase A (KGA) and glutaminase C (GAC). KGA protein levels were higher in male cells, but the difference was statistically insignificant. GAC protein levels were significantly higher in male cells (Fig. 4D). This is important, as GAC might play a more tumorigenic role than KGA (van den Heuvel et al., 2012; L. Xu et al., 2021). To determine whether greater GLS1 expression was coupled to greater dependency on its function, we treated male and female transformed astrocytes with varying doses of the GLS1 inhibitor CB-839. Notably, male cells displayed a steep dose-dependent sensitivity to GLS1 inhibition, whereas female cells were almost entirely resistant (Fig. 4E). This was true despite similar reductions in glutamate abundance to less than 50% of vehicle treated cells in both sexes, indicating no significant effect of sex on CB-839 potency (Fig. S3C). We confirmed sex differences in GLS1 dependency using BPTES, a less potent analog to CB-839, which also reduced male, but not female, cell number (Fig. S3D). To further validate sex-specific responses to GLS1 inhibition, we examined publicly available BPTES IC_50_ values in male and female tumor lines, spanning multiple cancer types (Daemen et al., 2018). Although female cell lines exhibited slightly higher IC_50_ values, these were not significantly different from the male lines (Fig. S3E). We then correlated BPTES IC_50_ values with GLS1 mRNA expression (Yang et al., 2019) and found that only male tumor lines exhibited a significant, but modest negative correlation between GLS1 expression and BPTES IC_50_ values (Fig. 4F), indicating that high GLS1 expression is associated with greater sensitivity to GLS1 inhibition in males, but not females.

The clinical and laboratory data thus far indicate that male glioma express higher levels of GLS1 and are more sensitive to GLS1 inhibition, suggesting they are more dependent on glutamine-derived glutamate. In addition, GLS1 expression is associated with BPTES sensitivity in male, but not female tumor cell lines, suggesting that sex differences in GLS1 dependency may extend beyond gliomas.

### Male transformed astrocytes are more dependent on glutathione for redox balance

To determine which metabolic pathways underlie the male-specific sensitivity to GLS1 inhibition, we assessed the effects of CB-839 on three metabolite groups linked with glutamine metabolism: TCA cycle, amino acids, and glutathione (Fig. 5A). While male and female cells both showed a reduction in TCA cycle metabolites upon GLS1 inhibition, the reduction was more profound in males. CB-839 treated cells exhibited heterogeneous responses in amino acid levels. Aspartate, glutamate, and proline, like TCA cycle metabolites, were more substantially reduced in male cells. In contrast, alanine, serine, asparagine, and threonine, were more substantially reduced in female cells. The remaining amino acids were increased in male and decreased in female cells. CB-839 treatment resulted in a significantly different (FDR<0.05) change in glutathione levels in male versus female cells. Males showed increased, and females showed decreased levels of glutathione, paralleling two of its amino acid components, glycine and cysteine. Thus, in male and female transformed astrocytes GLS1 inhibition has different metabolic effects.

**Figure 5:**
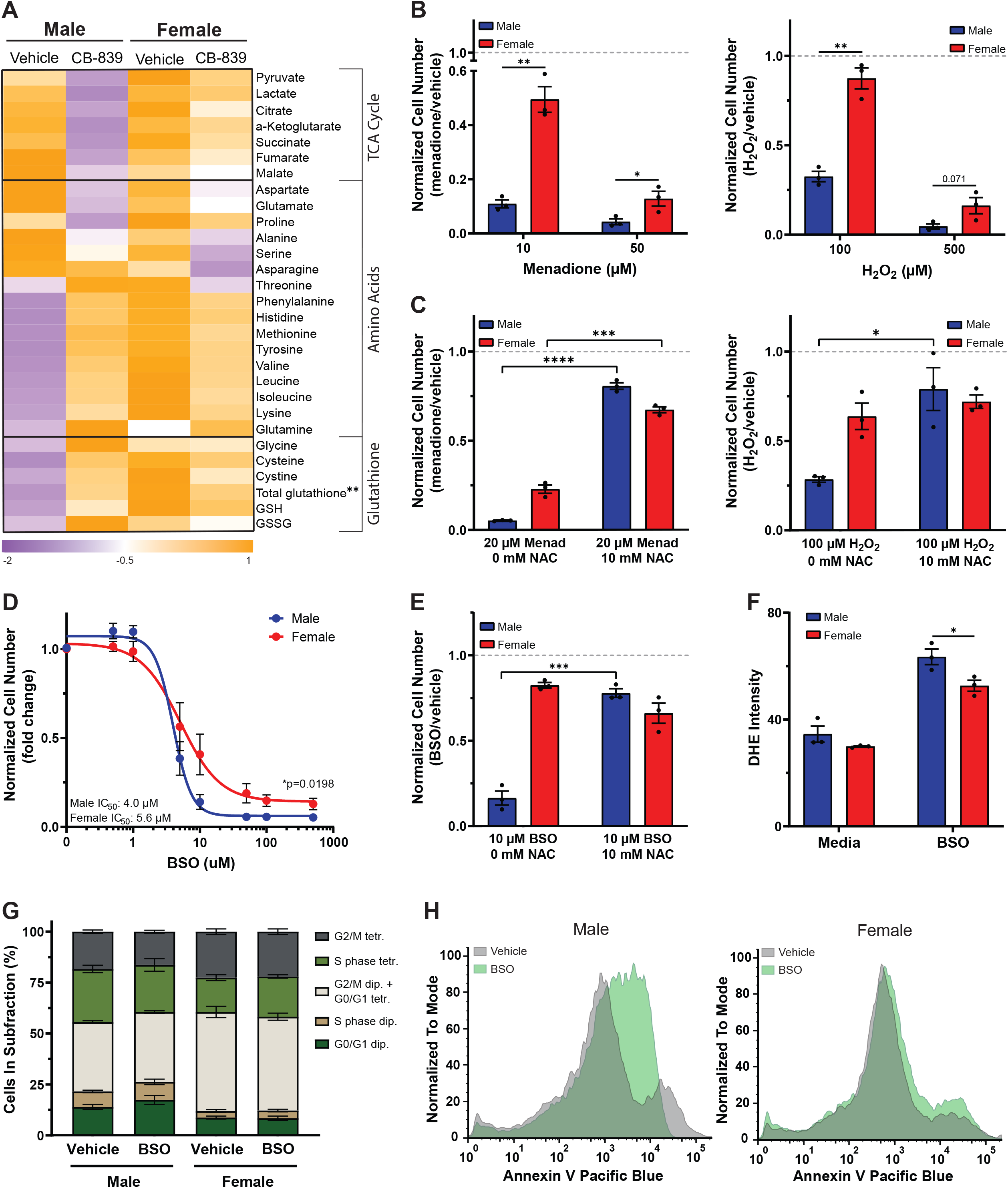
Male transformed astrocytes are more dependent on glutathione for redox balance. **A,** Heatmap of metabolite levels in transformed astrocytes treated with CB-839 or vehicle for 24 hrs. TCA cycle and amino acid metabolite abundance was measured via GC/MS; glutathione levels were measured using a spectrophotometric recycling assay. **B**, Cell number assay of transformed astrocytes treated with menadione or vehicle (left) or H_2_O_2_ or vehicle (right). **C**, Cell number assay of transformed astrocytes treated with menadione or vehicle and N-acetyl cysteine (NAC) (left) or H_2_O_2_ or vehicle and NAC (right). **D**, Buthionine sulfoximine (BSO) dose-response curves of transformed astrocytes. **E**, Cell number assay of transformed astrocytes treated with BSO or vehicle and NAC. **F**, Quantification of dihydroethidium (DHE) fluorescence intensity per cell in transformed astrocytes treated with BSO or vehicle. **G**, Quantification of propidium iodide histograms (Fig. S4D) of transformed astrocytes treated with BSO or vehicle. **H**, Representative Annexin V spectra of transformed astrocytes treated with BSO or vehicle. Spectra shown are representative spectra chosen from three independent. Data are mean +/- SEM of n=3/sex (three independent experiments, one male and female cell line), or for (D) mean +/- SEM of n=9/sex (three independent experiments, three separate male and female cell lines). **q<0.01; t-test, FDR adjusted p-values (A, G). *p<0.05, **p<0.01, ***p<0.001, ****p<0.0001; t-test (B-C, E-F). *p=0.0198; two-way ANOVA (D).

The greater reduction in TCA cycle metabolites suggests that male cells may be more dependent on glutamine for TCA cycle replenishment than female cells. The increase in glutathione levels and its amino acid components, further suggests that glutathione might play a more important role in the male metabolic stress response induced by GLS1 inhibition. Notably, male cells exhibited lower levels of glutathione and its amino acid components at baseline, suggesting that male cells may be more sensitive to oxidative stress. Sex differences in glutathione and ROS regulation have been described (Wang et al., 2020). Thus, we hypothesized that ROS regulation differs in male and female transformed astrocytes and that GLS1 inhibition has sex-specific effects on cellular redox balance.

To test our hypothesis, we first assessed whether male and female transformed astrocytes differ in their sensitivity to oxidative stress. We induced oxidative stress with menadione or hydrogen peroxide (H_2_O_2_) and found that both treatments resulted in a dose-dependent and significantly greater reduction in cell number in male versus female cells (Fig. 5B). Adding the thiol antioxidant N-acetylcysteine (NAC), significantly restored cell numbers in menadione treated male and female cells, and in H_2_O_2_ treated male cells (Fig. 5C), indicating that ROS accumulation and thiol oxidation are the major underlying reasons for the reduction in cell number upon menadione and H_2_O_2_ treatment. This suggests that male transformed astrocytes are more sensitive to oxidative stress than females.

To assess whether male cells are more dependent on glutathione to regulate redox balance, we depleted glutathione using buthionine sulfoximine (BSO), an inhibitor of glutamate cysteine ligase. Male cells were significantly more sensitive to glutathione depletion than female cells (Fig. 5D). This was not due to differences in glutathione levels upon treatment, as BSO treatment almost completely depleted glutathione in both sexes (Fig. S4A). This indicates that although female transformed astrocytes have significantly higher total glutathione levels at baseline, their greater resistance to BSO is not due to a greater abundance of cellular glutathione. We utilized a second strategy to deplete glutathione by inhibiting the glutamate/cystine antiporter X_CT_ (Cobler et al., 2018) with two X_CT_ inhibitors, sulfasalazine (SAS) and erastin and found that male cells were significantly more sensitive to both inhibitors (Fig. S4B-C).

To determine whether glutathione depletion affected cell number as a consequence of ROS accumulation and thiol oxidation, we treated male and female transformed astrocytes with vehicle or BSO with or without NAC. NAC significantly restored cell numbers in male and female BSO treated cells (Fig. 5E), and BSO induced significantly more dihydroethidium (DHE) oxidation – a measurement for superoxide accumulation – in males compared to females (Fig. 5F), suggesting increased levels of ROS. Thus, thiol oxidation may be the basis for cell number reduction in BSO treated cells. Lastly, we assessed whether BSO reduces cell number through cell cycle arrest or apoptosis. Neither male nor female cells underwent cell cycle arrest upon BSO treatment (Fig. 5G). However, in male cells, BSO increased Annexin V staining (Fig. 5H), indicating greater apoptosis. These data show that male transformed astrocytes are more dependent on glutathione to maintain their cellular redox balance, which might make them more dependent on GLS1 for the glutamate required in glutathione synthesis.

### Male transformed astrocytes require glutamine to replenish their TCA cycle

To assess whether sex-specific effects of GLS1 inhibition on cell number are driven by a cellular redox imbalance, we measured DHE oxidation in CB-839 treated male and female transformed astrocytes. Similar to glutathione depletion, CB-839 resulted in a significantly greater DHE oxidation in male cells (Fig. 6A). However, in contrast to BSO, the CB-839 growth phenotype was not rescued by NAC (Fig. 6B). Thus, thiol oxidation is not the underlying reason for the male-specific sensitivity to CB-839. To gain greater insight, we assessed the effects of GLS1 inhibition on the cell cycle and apoptosis. Unlike their response to glutathione depletion, male transformed astrocytes underwent cell cycle arrest in response to GLS1 inhibition (Fig. 6C) without an increase in apoptosis (Fig. 6D). Thus, the mechanisms underlying changes in cell number are different upon glutathione depletion (apoptosis) and GLS1 inhibition (cell cycle arrest), indicating that while male cells are more dependent on glutathione, redox balance is not the basis for their greater sensitivity to GLS1 inhibition.

**Figure 6:**
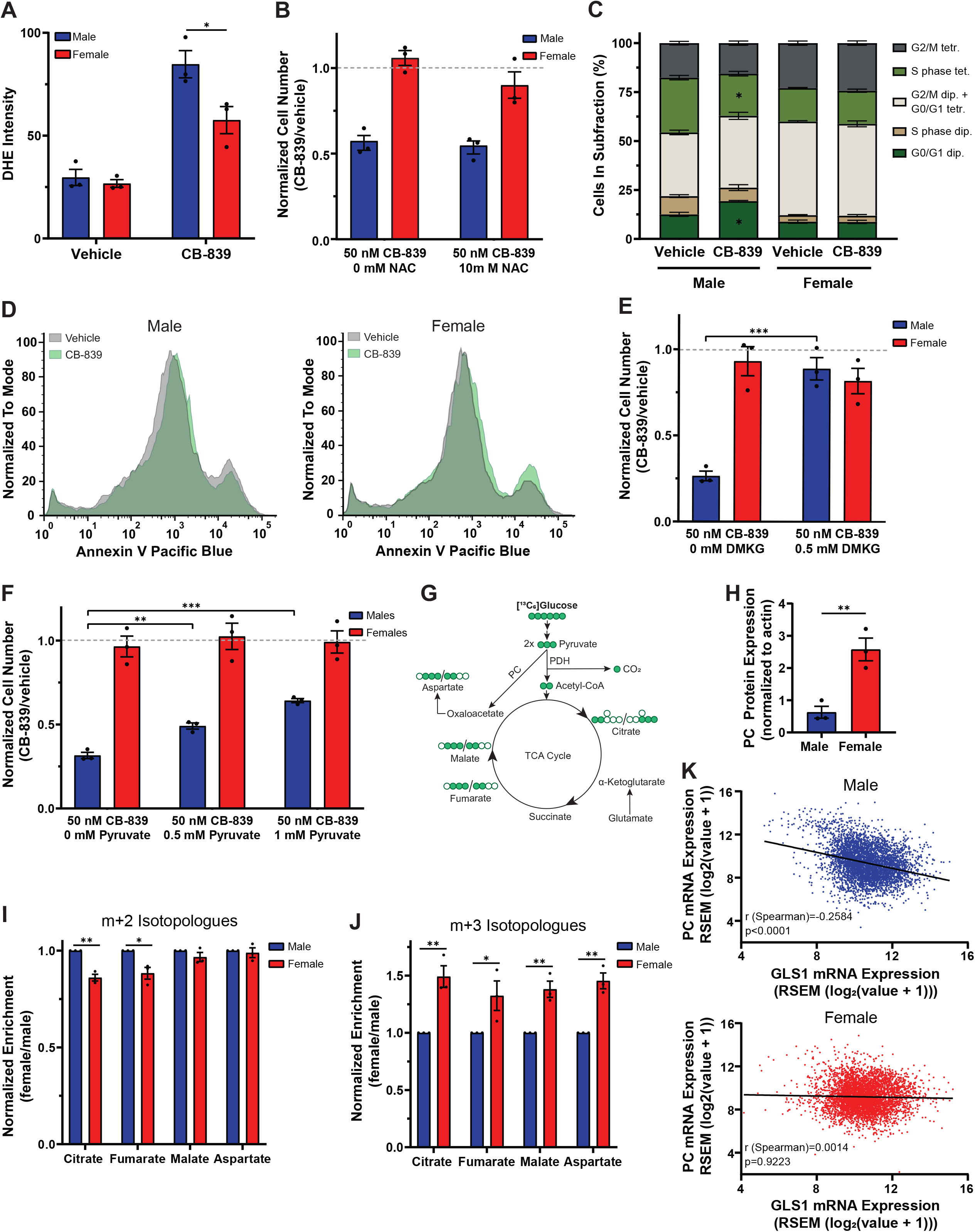
Male transformed astrocytes require glutamine to replenish their TCA cycle. **A**, Quantification of dihydroethidium (DHE) fluorescence intensity per cell in transformed astrocytes treated with CB-839 or vehicle. **B**, Cell number assay of transformed astrocytes treated with CB-839 or vehicle and N-acetyl cysteine (NAC). **C**, Quantification of propidium iodide histograms (Fig. S5C) of transformed astrocytes treated with CB-839 or vehicle. **D**, Representative Annexin V spectra of transformed astrocytes treated with CB-839 or vehicle. Spectra shown are representative spectra chosen from three independent experiments. **E**, Cell number assay of transformed astrocytes treated with CB-839 or vehicle and dimethyl-α-ketoglutarate (DMKG). **F**, Cell number assay of transformed astrocytes treated with CB-839 or vehicle and pyruvate. **G**, TCA cycle flux schematic. **H**, Western blot quantification of PC. Expression values are normalized to β-actin. **I-J,** Label incorporation of [^13^C_6_]glucose in transformed astrocytes. Label incorporation into metabolites was measured via GC/MS. m+2 **(I)** and m+3 isotopologues **(J)** after one turn of the TCA cycle are shown. **K,** Nonparametric Spearman correlation between PC and GLS1 mRNA expression levels in n=4714 male and n=4982 female pan-cancer patient samples. Data are mean +/- SEM of n=3/sex (three independent experiments, one male and female cell line). *p<0.05, **p<0.01, ***p<0.001; t-test (A, B, E, F, H). *q<0.05, **q<0.01; t-test, FDR adjusted p-values (C, I-J).

Glutamate is also required for α-KG synthesis, which is important in multiple cellular processes, including replenishing the TCA cycle. Since GLS1 inhibition with CB-839 resulted in a greater reduction in TCA cycle metabolites in male transformed astrocytes, we hypothesized that TCA cycle metabolite depletion triggers growth arrest in male cells. To test this hypothesis, we attempted to rescue CB-839 treated male and female transformed astrocytes with two metabolites commonly used for TCA cycle restoration, dimethyl-α-KG (DMKG) and pyruvate. DMKG fully rescued the male growth phenotype (Fig. 6E). Pyruvate had a weaker, but dose-dependent and significant rescue effect (Fig. 6F). DMKG is a TCA cycle metabolite and therefore a more direct rescue of TCA cycle function compared to pyruvate, which may explain the greater potency of DMKG. Thus, male transformed astrocytes are sensitive to GLS1 inhibition due to its inhibitory effects on TCA cycle activity.

TCA cycle metabolites can also be replenished by glucose-derived pyruvate, which can enter the TCA cycle via two enzymatic reactions. Most pyruvate entering the TCA cycle is converted to acetyl-CoA by pyruvate dehydrogenase (PDH). However, pyruvate carboxylase (PC) can also directly convert pyruvate to oxaloacetate (Fig. 6G). PC activity can support glutamine-independent cancerous growth as it provides a compensatory anaplerotic mechanism (Cheng et al. 2011). We assessed PC protein expression in transformed astrocytes and found that females have ≈4-fold higher expression of PC (Fig. 6H and S5A), which was unaffected by CB-839 treatment (Fig. S5B). Next, we assessed PC activity in male and female cells at baseline. We incubated cells in [^13^C_6_]glucose and assessed label enrichment. Upon its conversion from [^13^C_6_]glucose, [^13^C_3_]pyruvate can replenish the TCA cycle via PDH or PC. PDH transfers two, and PC three, labeled carbons into the TCA cycle, producing either m+2 or m+3 isotopologues on the cycle’s first turn (Fig. 6G) (Chen et al., 2019). We measured m+2 and m+3 isotopologues of aspartate, malate, fumarate, and citrate. Label enrichment in citrate and fumarate m+2 isotopologues was higher in males, suggesting a slightly higher utilization of PDH for glucose input into the TCA cycle in male cells (Fig. 6I). In contrast, females exhibited ≈1.5-fold greater label incorporation into all m+3 isotopologues (Fig. 6J), indicating they have greater PC capacity for feeding glucose into the TCA cycle. Thus, male cells are more reliant on glutamate for TCA cycle replenishment. Interestingly, GLS1 and PC mRNA expression levels in the TCGA Pan-Cancer Atlas were negatively correlated in male, but not female pan-cancer samples (Fig. 6K). This finding suggests that male cancers with higher GLS1 expression and therefore greater sensitivity to CB-839 would have the least amount of PC-mediated resistance. Thus, we conclude that female cancer cells possess greater metabolic flexibility for TCA cycle replenishment, resulting in resistance to therapeutic targeting of glutamine metabolism.

## Discussion

Metabolic reprogramming supports the high energetic and biosynthetic demands of cancerous growth. Targeting cancer’s unique metabolic adaptations holds great promise as a treatment with a high therapeutic index. For this approach to work optimally for all patients, it must take sex differences in energy and biosynthetic pathways into account. Here, we report for the first time that cancer cells retain normal sex differences in glutamine metabolism. Sex differences in glutamine uptake, glutamine metabolism, and GLS1 expression were evident in our mouse transformed astrocyte cell lines, human glioma cell lines, and GBM patient samples, as well as in normal brain, suggesting that sex differences in glutamine metabolism are conserved across species and between cancerous and non-cancerous tissue. Similar sex differences in glutamate levels and glutamine metabolism have been reported in normal brain tissues of rats (Al-Suwailem et al., 2018; Davis et al., 1999).

We focused on glutamine metabolism because it is being targeted in early phase cancer clinical trials (NCT02071927, NCT04250545, NCT03528642, NCT03965845). We found that male cells were significantly more dependent on glutamine, which they shuttled into all its major downstream processes. GLS1 inhibition resulted in greater growth arrest in male cells, and sex-dependent effects on the TCA cycle and glutathione metabolism, which played a greater role in male redox balance. The underlying reason for sex differences in sensitivity to GLS1 inhibition was PC-mediated TCA cycle replenishment by glucose, which was more active in female cells.

In normal metabolism, sex differences are evident throughout life (Hedrington & Davis, 2015; Lamont et al., 2003; Ray et al., 1995; Tiffin et al., 1991). Here, we show for the first time that they are also a feature of cancer. There was substantial concordance between the male-biased metabolic pathways we identified in GBM, and those described in male serum (Krumsiek et al., 2015). For example, amino acid and carbohydrate metabolites were increased in males, while females were enriched in certain lipids in both datasets. While these findings were highly significant in serum, they did not reach significance in the GBM dataset, which is most likely due to its small sample size. Overall, these findings suggest that fundamental sex differences in metabolism are present in GBM and may even extend beyond gliomas.

As glutamine metabolism differs in male and female cells, response to targeting glutamine metabolism might need to be adapted to sex-biased mechanisms of resistance. Male glioma patients may respond better to CB-839 treatment than female glioma patients and co-targeting PC activity might block resistance to GLS1 in female cells.

We also showed that redox regulation differs in male and female transformed astrocytes. Sex differences in sensitivity to ROS, pro-oxidant levels, and ROS regulation are well described in healthy brain tissue (Rubin et al., 2020). Here, we provide evidence that these sex differences might persist in cancerous tissue and could result in sex-specific responses to treatments that depend on ROS accumulation. This study is not the first study providing evidence that novel anti-cancer therapeutics currently in clinical trials might have sex-specific effects. It was recently reported that clinically utilized bromodomain inhibitors could produce sex-specific responses (Kfoury et al., 2021). Together, these studies emphasize the importance of including sex as a variable in basic research and clinical settings.

Consistent sex differences in metabolite abundance in cancerous and non-cancerous tissue lead us to hypothesize that sex differences in glutamine metabolism might be conserved across other cancers, a possibility that should be explored for optimized clinical use of GLS1 inhibition. Sex differences in other metabolic pathways may be similarly important. As an example, all branched-chain amino acids (BCAAs) were included in the top 75 enriched metabolite list and exhibited a male-bias. Our glutamine tracing experiments showed that male transformed astrocytes incorporated ≈2-fold more label into BCAAs, indicating that targeting BCAA metabolism may be a unique vulnerability in male cells. Finally, our data suggest that other metabolic pathways, such as carbohydrates and fatty acids, exhibit sex differences in GBM. It remains to be determined how sex differences in these metabolic groups affect cancer biology and treatment response.

### Limitations of the study

While the sex-specific analysis of human GBM metabolite abundance shows remarkable similarities with previously published studies, it is underpowered to detect significant sex differences. A greater number of patient samples would be necessary to validate those findings.

Here, we show that male transformed astrocytes are dependent on glutamine for their growth because they depend on α-KG for TCA cycle replenishment. However, α-KG plays additional metabolic and epigenetic roles. It is an essential co-factor for Jumonji-domain containing histone demethylases and Ten Eleven Translocation proteins which mediate histone and DNA demethylation, respectively. It is also an important component of transamination and amino acid synthesis. These cellular processes are likely to also be affected by GLS1 inhibition. Thus, it remains to be determined whether the disruption of other cellular processes downstream of α-KG also contributes to the observed growth arrest in male cells following GLS1 inhibition.

All mechanistic *in vitro* studies were performed in the same model of transformed astrocytes. While we repeated several key experiments (glutamine deprivation, CB-839 treatment, BSO treatment) in three separately established male and female transformed astrocyte cell lines, the translational relevance of the work could be improved by using additional *in vitro* models. Despite this limitation, we believe that the validation of our *in vitro* data using human samples and cell line data throughout the paper is strong and suggests that sex differences in GLS1 dependency are not likely to be a model-specific phenotype. Lastly, *in vivo* studies including human xenograft models and spontaneous tumor models should be conducted to confirm and extend our findings to other types of glioma and other cancer models.

## Acknowledgements

This work was supported by the Cancer Biology Pathway Molecular Oncology Training Grant NIH T32CA113275 (JSp), the NIH grants R01 CA174737-06 (JBR), K99/R00 CA218869 (JEI), R21 CA242221 (JEI), R35 ES028365 (GJP), P01 CA217797 (DRS), Joshua’s Great Things (JBR), the Alvin J. Siteman Cancer Center Investment Program / The Foundation for Barnes-Jewish Hospital (JBR, JEI) and the Barnard Research Fund (JBR, JEI). SK was supported in part by the NIH/NCI Paul Calabresi Career Development Award for Clinical Oncology K12 CA184746. MSK core facilities are supported by NIH/NCI Cancer Center Support Grant P30 CA008748.

The results shown here are in part based upon data generated by the TCGA Research Network: https://www.cancer.gov/tcga. We thank the Alvin J. Siteman Cancer Center at Washington University School of Medicine and Barnes-Jewish Hospital in St. Louis for the use of the Siteman Flow Cytometry Core. The Siteman Cancer Center is supported in part by an NCI Cancer Center Support Grant P30 CA091842. In particular, we thank William Eades for his help with flow cytometry. We thank the Biomedical Mass Spectrometry Core at Washington University School of Medicine for providing GC/MS equipment. The core is supported by the NIH grant R24 GM136766. We thank Michael McCormick and the Radiation and Free Radical Research Core at the University of Iowa for their help with assessing cellular GSH and GSSG levels.

## Author Contributions

Conceptualization: JSp, JBR, and JEI. Investigation and formal analysis: JSp, JKJ, and SZ performed and analyzed most *in vitro* experiments; EN, HG, and OT helped with data collection and analysis; CF collected and analyzed GC/MS data; KC collected and analyzed LC/MS data; JSc collected and analyzed solid-state NMR data; MSG, AM, MG, MD, and SK collected and analyzed [^18^F]FGln data, JL analyzed GBM metabolite data. Data curation: JL. Supervision: KLT, DRS, GJP, JL, JBR, and JEI. Resources: PC. Visualization: JSp. Writing – original draft: JSp. Writing – review & editing: JBR, and JEI, as well as all authors. Funding acquisition: JBR, and JEI.

## Methods

### Resource Availability

#### Lead Contact

Further information and request for resources and reagents should be directed to and will be fulfilled by the lead contact Joseph E. Ippolito.

#### Materials Availability

There are no restrictions to the availability of all materials mentioned in the manuscript. Data and Code Availability

Original data for Figure 1 in the paper is available in (Prabhu et al., 2019). The tumor cell line dataset used in Figure 4H in the paper is available in (Daemen et al., 2018). Targeted GC/MS metabolite data and isotopologue data generated for this publication are included in the published article.

### Experimental Model and Subject Details

#### Cell culture

The male and female *Nf1-/-DNp53* astrocytes (transformed astrocytes) were previously generated and published by our lab (Sun et al., 2014). Briefly, primary cultures of male and female *Nf1-/-* astrocytes were isolated from the neocortices of individual postnatal day 1 *Nf1fl/fl GFAP-Cre* mice as described previously (Dasgupta, 2005; Sun et al., 2014) and cultured in Gibco Dulbecco’s Modified Eagle Medium: Nutrient Mixture F-12 (DMEM/F-12), supplemented with 10% fetal bovine serum (FBS), and 1% penicillin/streptomycin (PS) at 37°C and 5% CO_2_.The sex of each mouse was determined from tail DNA by PCR for *Jarid1c* and *Jarid1d* (Clapcote & Roder, 2005; J. Xu et al., 2008). Cells from at least three male and three female mice across multiple litters were pooled. Male and female *Nf1−/−* astrocytes were then infected with retrovirus encoding a flag-tagged dominant-negative form of p53 (DNp53) and EGFP as previously described (Sun et al., 2014). The DNp53 plasmid consists of amino acids 1–14 of the transactivation domain followed by amino acids 303–393, thus lacking the DNA binding domain. These male and female *Nf1-/-; DNp53* are referred to as male and female transformed astrocytes. Multiple astrocyte harvests and retroviral infections were performed, which led to multiple lines of male and female GBM astrocytes, which are referred to as “lots” of male and female transformed astrocytes. Experiments were performed with up to three different lots of transformed astrocytes. All transformed astrocytes were cultured in DMEM/F-12, supplemented with 10% FBS at 37°C and 5% CO_2_. All experiments were performed at passages 5 to 12.

#### Patient samples

As part of a prospective research protocol approved by the Institutional Review Board, patients with glioma were imaged with [^18^F]FGln PET/CT. All procedures were performed in accordance with the ethical standards of the institutional and/or national research committee and with the 1964 Helsinki declaration and its later amendments or comparable ethical standards. All patients provided written informed consent [NCT 01697930].

### Method Details

#### Human GBM metabolite analysis

The male and female human GBM metabolite dataset (Prabhu et al., 2019) and the sex of each surgical specimen was kindly provided by Dr. Prakash Chinnaiyan. The dataset included 44 male and 32 female GBM surgical specimens and 757 metabolites. Upon initial normalization described previously (Prabhu et al., 2019), metabolites showing no variation (same expression across all samples) were removed, leaving 718 metabolites for further analysis. Then, the data was log_2_ transformed and the z-score values were derived for each metabolite. The average z-score across metabolites belonging to each sub- or super-pathway was calculated to represent the metabolic pathway activity of the sub- or super-pathway. The R/Bioconductor software package limma version 3.48.3 (Ritchie et al., 2015) was used for sex-differential analysis at the individual metabolite level, the sub-pathway level, and the super-pathway level separately.

#### Heatmap

Z-score values from each metabolite were represented in a heatmap using Morpheus (https://software.broadinstitute.org/morpheus). Hierarchical clustering was performed using Morpheus and samples were clustered based on Euclidean distance and average linkage. K-means clustering was performed using Morpheus and samples were clustered based on Euclidean distance.

#### Glutamine tracer PET imaging

##### [^18^F]fluoroglutamine (FGln) PET/CT imaging protocol

[^18^F]FGln was synthesized by Memorial Sloan Kettering Cancer Center’s Radiochemistry and Molecular Imaging Probe Core Facility as described (Dunphy et al., 2018). Each [^18^F]FGln dose met drug product acceptance specifications, including radiochemical purity and identity, residual solvent content, endotoxin content, radionuclidic identity, pH, and appearance. The median administered activity was 241.98 MBq (6.54 mCi); IQR: 22.2 MBq (0.60 mCi). The tracer was given as a rapid intravenous bolus injection. Static PET scans were performed at 78 min ± 9 min after injection (15 to 30 min acquisition time) on a Discovery STE, 690, or 710 camera (GE Healthcare). A CT scan (120 kVp, 70 mA, and 3.8 mm slice thickness) was obtained for attenuation correction, anatomic localization, and coregistration purposes. PET emission data were acquired in 3-dimensional mode; corrected for attenuation, scatter, and random events; and iteratively reconstructed into a 256 × 256 × 47 matrix (voxel dimensions, 1.95 × 1.95 × 3.27 mm) for brain lesions using the ordered-subset expectation maximization algorithm provided by the manufacturer.

##### [^18^F]FGln PET image interpretation and data analysis

A board-certified nuclear medicine physician defined regions of interest (ROIs) for the lesions and normal brain on a GE Advantage workstation using PET VCAR software. The [^18^F]FGln PET/CT was windowed to visualize the focal [^18^F]FGln avidity associated with the known brain lesion on MRI then an ROI was placed to encompass the entire area of abnormal [^18^F]FGln avidity. For normal brain tracer uptake in the grey matter, an ROI was placed contralaterally to the brain lesion. Lesion volumes were measured by performing three-dimensional threshold-based volume of interest (VOI) analyses for the [^18^F]FGln uptake in all patients. Tracer uptake was quantified by SUVs and corrected for lean body mass (SUL) using the sex-specific Janmahasatian formulation (Tahari et al., 2014). For lesion VOI, SUL_max_ was recorded. SUL_max_ referred to the voxel with the maximum intensity of [^18^F]FGln uptake in the VOI. For patients with multiple lesions, we used the lesion with the highest SUL_max_. For brain VOI, SUL_mean_ was recorded. SUL_mean_ referred to the mean intensity of [^18^F]FGln uptake in the VOI. Representative axial images were chosen from a male and from a female high-grade glioma patient (IDH WT, MGMT unmethylated, recurrent tumor biopsy confirmed 12 days and within 1 month, respectively, after scan). Axial contrast-enhanced T1-weighted MRI (T1W + C MRI) axial images, PET and fused PET/MRI images are shown.

#### Drug preparation

CB-839 was purchased as a powder and diluted in DMSO to a stock concentration of 10 mM. BPTES was purchased as a powder and diluted in DMSO to a stock concentration of 100 mM. Menadione was purchased as a powder and diluted in DMSO to a stock concentration of 150 mM. Buthionine sulfoximine (BSO) was purchased as a powder and diluted in water to a stock concentration of 100 mM.

#### Cell number assays

##### Glutamine dose-response curve

Cell number was assessed using the colorimetric sulforhodamine B assay (Vichai & Kirtikara, 2006). Briefly, cells were plated in triplicates in a 96-well plate at a density of 1,000 cells/well in DMEM/F-12 media supplemented with 10% FBS. For each experiment, a standard curve for male and female cells was plated at the same time. 6 hrs after plating, standard curves were fixed as described below. 24 hrs after plating, media was changed to DMEM/F-12 (USBiological) containing 1.2 g/L sodium bicarbonate, 10 mM HEPES, 10% dialyzed FBS, 10 mM glucose, and glutamine at various concentrations (4.5, 2.5, 1.5, 1, 0.5, 0.1, 0 mM). Cells were incubated for 4 days. On day 4, plates were fixed with ice cold 5% trichloroacetic acid at 4°C for 1 hr. Then, plates were washed 4 times with tap water and allowed to dry at room temperature overnight. Next, plates were stained with 0.057% (wt/vol) sulforhodamine B (SRB) in 1% (vol/vol) acetic acid at room temperature for 30 min. The plates were washed quickly 3 times with 1% (vol/vol) acetic acid and allowed to dry at room temperature overnight. SRB was dissolved in 10 mM unbuffered Tris-Base solution (pH 10.5) on a shaker at room temperature for at least 30 min. Absorbance was measured at 510 nm using a Synergy HT microplate reader (BioTek). Absorbance values were converted to interpolated cell numbers in GraphPad Prism, using the standard curve absorbance values for each experiment.

##### CB-839, buthionine sulfoximine (BSO), H_2_O_2_, and menadione treatment

Cell number was assessed as described above with minor modifications. Cells were plated in triplicates in a 96-well plate at a density of 2,000 cells/well in DMEM/F-12 media supplemented with 10% FBS. 24 hrs after plating, cells were treated with various doses of CB-839 (0.01, 0.025, 0.05, 0.1, 0.5, 1, 5 µM) or vehicle (DMSO), BSO (0.5, 1, 5, 10, 50, 100, 500 µM) or vehicle (water), H_2_O_2_ (100 µM and 500 µM) or vehicle (water), or menadione (10 µM and 50 µM) or vehicle (DMSO). Both male and female cells were around 50% confluent when treatment was started. Cells were incubated for 48 hrs.

##### Rescue assays

Cell number was assessed as described above with minor modifications. Cells were plated in triplicates in a 96-well plate at a density of 2,000 cells/well in DMEM/F-12 media supplemented with 10% FBS. 24 hrs after plating, cells were treated with either CB-839 (50 nM) or vehicle (DMSO), menadione (20 µM) or vehicle (DMSO), or H_2_O_2_ (100 µM) or vehicle (water) and either N-acetyl-cysteine (NAC) (10 mM), or dimethyl-α-ketoglutarate (DMKG) (0.5 mM), or pyruvate (0.5 mM and 1 mM). Both male and female cells were around 50% confluent when treatment was started. Cells were incubated for 48 hrs.

A 1 M NAC stock solution was prepared fresh before each experiment. 188 mg NaHCO3 and 326 mg NAC were dissolved in 2 mL molecular grade water. pH was adjusted to 7.0 and solution was filter sterilized with a 70 µm filter. An 800 mM pyruvate stock solution in media was prepared fresh and filter sterilized with a 70 µm filter before each experiment. DMKG was directly added to the media.

#### Glutamine consumption assay

##### Sample preparation

Cells were plated in a 6-well plate at a density of 50,000 cells/well in DMEM/F-12 media supplemented with 10% FBS. 6 technical replicates were plated. 24 hrs after plating, media was changed to DMEM/F-12 (USBiological) containing 1.2 g/L sodium bicarbonate, 10 mM HEPES, 10% dialyzed FBS, 10 mM glucose, and 4.5 mM glutamine. Cells were incubated for another 24 hrs. Then, media was collected from each well. Next, 20 µL of each media sample was added to 180 µL of a 2:2:1 methanol:acetonitrile:water (−20°C) mixture. Then, samples were vortexed for 1 min and placed at −20°C for 1 hr. Next, samples were centrifuged at 16,000 g at 4°C for 10 min and supernatant was transferred to LC glass vials. Samples were stored at −80°C until LC/MS analysis (see below).

#### Stable isotope tracing with LC/MS

##### Sample preparation

Cells were plated in 100 mm dishes at a density of 250,000 cells/well in DMEM/F-12 media supplemented with 10% FBS. 6 technical replicates were plated. 24 hrs after plating, media was changed to DMEM/F-12 (USBiological) containing 1.2 g/L sodium bicarbonate, 10 mM HEPES, 10% dialyzed FBS, 10 mM glucose, and 4.5 mM glutamine or [^13^C_5_ N_2_]glutamine. Cells were incubated for another 24 hrs. Then metabolites were extracted at 75-85% confluence. For metabolite extraction, media was removed, cells were washed on dry ice 3 times with ice-cold 1X PBS, followed by 3 washing steps with water. Then, cells were scraped into methanol (−20°C) and transferred to Eppendorf tubes. Next, samples were dried at room temperature using a Labconco CentriVap Concentrator. Cells were lysed by adding a 2:2:1 methanol:acetonitrile:water (−20°C) mixture to the dried pellet. Then, cells were vortexed for 30 sec, placed into a liquid N2 bath for 1 min, and bath sonicated at 25°C for 10 min. This process was repeated 3 times. Then, samples were stored at −20°C overnight. The next day, samples were centrifuged at 12,000 g at 4°C for 10 min. The supernatant was transferred to a new tube and dried at room temperature using a Labconco CentriVap Concentrator. Total protein content of pellets was measured by Pierce BCA protein assay kit, following the manufacturer’s microplate assay protocol.

Then, 1 µL of a 2:1 acetonitrile:water (−20°C) mixture was added to the dried pellet per 2.5 µg of protein. Next, the samples were bath sonicated at 25°C for 5 min and vortexed for 1 min. This process was repeated 2 times. Then, samples were stored at 4°C overnight. The next day, samples were centrifuged at 16,000 g at 4°C for 10 min and the supernatant was transferred to LC glass vials and stored at −80°C until LC/MS analysis.

##### LC/MS analysis

Ultra-high performance LC (UHPLC)/MS was performed with a Thermo Fisher Scientific Vanquish Horizon UHPLC system interfaced with a Thermo Fisher Scientific Orbitrap ID-X Tribrid Mass Spectrometer. Hydrophilic interaction liquid chromatography (HILIC) was conducted with a HILICON iHILIC-(P) Classic guard column (20 mm x 2.1 mm, 5 µm) connected to a HILICON iHILIC-(P) Classic HILIC column (100 mm x 2.1 mm, 5 µm). Mobile-phase solvents were composed of A=20 mM ammonium bicarbonate, 0.1% ammonium hydroxide and 2.5 µM medronic acid in water:acetonitrile (95:5) and B=2.5 µM medronic acid in acetonitrile:water (95:5). The column compartment was maintained at 45°C for all experiments. The following linear gradient was applied at a flow rate of 250 µL min-1: 0-1 min: 90% B, 1-12 min: 90-35% B, 12-12.5 min: 35-25% B, 12.5-14.5 min: 25% B. The column was re-equilibrated with 20 column volumens of 90% B. The injection volume was 3 µL for cell samples and 4 µL for media samples.

Data were collected with the following settings: spray voltage, −3 kV; sheath gas, 35; auxiliary gas, 10; sweep gas, 1; ion transfer tube temperature, 250 °C; vaporizer temperature, 300 °C, mass range, 70-1000 Da, and two narrow-mass ranges, 270-400 Da and 550-700 Da; resolution, 240,000 (MS1), 30,000 (MS/MS); maximum injection time, 500 ms; isolation window, 1.5 Da; polarity, negative. LC/MS data were processed and analyzed with the open-source Skyline software (Adams et al., 2020; MacLean et al., 2010). Natural-abundance correction of ^13^C was performed with AccuCor (Su et al., 2017).

#### Stable isotope tracing with GC/MS

##### Sample preparation

Cells were plated in a 6-well plate at a density of 50,000 cells/well in DMEM/F-12 media supplemented with 10% FBS. 6 technical replicates were plated. For glutamine labeling experiments, media was changed to DMEM/F-12 (USBiological) containing 1.2 g/L sodium bicarbonate, 10 mM HEPES, 10% dialyzed FBS, 10 mM glucose, and 4.5 mM glutamine or [^13^C_5_ N_2_]glutamine 24 hrs after plating. For glucose labeling experiments, media was changed to DMEM/F-12 (USBiological) containing 1.2 g/L sodium bicarbonate, 10 mM HEPES, 10% dialyzed FBS, 4.5 mM glutamine, and 10 mM glucose or [^13^C_6_]glucose 24 hrs after plating. Cells were incubated for another 24 hrs. Then metabolites were extracted at 75-85% confluence. For metabolite extraction, media was removed, and cells were washed on dry ice 2 times with ice-cold 1X PBS. Then, cells were scraped into 200 µL of an 80:20 methanol:water (−20°C) mixture. Two wells were combined per one technical replicate. Next, samples were centrifuged at 12,000 g at 4°C for 20 min. The supernatant was transferred to a flat bottom GC insert vial and dried at room temperature using a Labconco CentriVap Concentrator. Then, the dried pellets were derivatized with 14 µL of 20 mg/mL methoxyamine HCl in pyridine and incubated on at 38°C for 90 min. Immediately following the 90 min incubation, 56 µL of MTBSTFA (with 1% t-BDMCS were added and samples were incubated at 38°C for 30 min (Fiehn, 2016).

##### GC/MS analysis

Derivatized samples were run on an Agilent 7890A GC coupled to an Agilent 5975C MS and data was acquired and analyzed in MSD ChemStation E.02.02.1431 (Agilent). The GC temperature program was set to: initial temperature: 60°C, fold for 0.65 min; temperature rate 1: 11.5°C/min to 201°C, hold for 0.65 min; temperature rate 2: 3.5°C/minute to 242°C, hold for 0 min; temperature rate 3: 16°C/min to 300°C, hold for 7 minutes. Data on all isotopologues was collected in SIM mode (see Table S2 for m/z monitored). Isotopologue peak areas in each sample were normalized in FluxFix (Trefely et al., 2016) to isotopologues of unlabeled samples to adjust for the natural abundance of each isotopologue.

#### Solid-state NMR

##### Sample preparation

Experimental design and data analysis and interpretation were consistent with previous publications (Chen et al., 2016; Yao et al., 2016). Cells were plated in 150 mm dishes at a density of 300,000 cells/dish in DMEM/F-12 media supplemented with 10% FBS. 24 hours after plating, media was changed to DMEM/F-12 (USBiological) containing 1.2 g/L sodium bicarbonate, 10 mM HEPES, 10% dialyzed FBS, 10 mM glucose, and 4.5 mM [^13^C_5_ N_2_]glutamine. Cells were incubated for 48 hrs. Then, cells were trypsinized and centrifuged at 200 g at 4°C for 5 min. Media was aspirated and the cell pellet was washed with ice cold 1X PBS. Washing was repeated two times and the cell pellet was stored at −80°C. Before NMR analysis the cell pellet was dried in a lyophilizer at −20°C. The total weight of the male and female dried cell pellet submitted for solid-state NMR analysis was 20 mg each and was accumulated over three sample preparations.

##### Solid-state NMR spectrometer and pulse sequences

Experiments were performed at 12 Tesla with a transmission-line probe having a 12-mm long, 6-mm inner-diameter analytical coil, and a Chemagnetics/Varian ceramic spinning module. Samples were spun using a thin-wall Chemagnetics/Varian (Fort Collins, CO/Palo Alto, CA) 5-mm outer diameter-zirconia rotor at 8000 Hz, with the speed under active control and maintained to within ±2 Hz. A Tecmag Libra pulse programmer (Houston, TX) controlled the spectrometer. Two-kW American Microwave Technology (AMT) power amplifiers were used to produce radio-frequency pulses for ^13^C (125.7 MHz) and ^15^N (50.3 MHz). The ^1^H (500 MHz) radio-frequency pulses were generated by a 2-kW Creative Electronics tube amplifier driven by a 50-W AMT amplifier. The *π*-pulse lengths were 8 μs for both ^13^C and ^1^H, and 9 µs for ^15^N. Proton-carbon-matched cross-polarization transfers were made in 2 ms at 56 kHz. Proton dipolar decoupling was 100 kHz during data acquisition. The ^13^C spectra were obtained with 80-μs interrupted decoupling which enhances signals from non-protonated carbonyl carbons (175 ppm), and from protonated carbons whose dipolar coupling to protons is reduced by molecular motion (30 ppm). The ^13^C chemical-shift scale was referenced to tetramethylsilane. The ^15^N chemical-shift scale was referenced to solid ammonium sulfate. (To switch to liquid ammonia as a secondary reference, add 20 ppm.) All of the ^13^C and ^15^N NMR signals were due to labeling. Natural abundance signals were negligible.

#### mRNA expression data

RNA-Sequencing and mRNA microarray data were downloaded from cBioPortal for cancer genomics (http://www.cbioportal.org/) (Cerami et al., 2012; Gao et al., 2013).

For glioma cell lines and tumor cell lines, mRNA expression data from the Cancer Cell Line Encyclopedia (Ghandi et al., 2019; Nusinow et al., 2020) were downloaded and graphed as mRNA expression (RNA Seq RPKM (log_2_(value + 1))) data. Only cell lines that had a sex assigned in the Cancer Cell Line Encyclopedia were downloaded. Then, the sex of the cell lines was confirmed using the Cellosaurus data resource (https://web.expasy.org/cellosaurus/). Cell lines with a discrepancy between the sex assigned by the Cancer Cell Line Encyclopedia and by the Cellosaurus data resource were excluded. For tumor cell lines, only cell lines with a measurable IC_50_ (Daemen et al., 2018) were included in the analysis.

For GBM patient samples, mRNA expression data from the TCGA Firehose Legacy GBM patient dataset (Brennan et al., 2013; Cancer Genome Atlas Research Network, 2008) were downloaded and graphed as mRNA expression (microarray) data. Only samples with a known sex and IDH wild-type status were included in the analysis. IDH status of each sample was determined using the IDH mutational status from the TCGA Firehose Legacy GBM patient dataset and the TCGA Merged Cohort of LGG and GBM dataset (Ceccarelli et al., 2016).

For pan-cancer patient samples, mRNA expression data from the TCGA Pan-Cancer Atlas (Hoadley et al., 2018) were downloaded and graphed as mRNA expression (RNA Seq RSEM (log_2_(value + 1))) data. Only samples with a known sex status were included in the analysis.

#### Western blotting

To extract protein, cells were harvested at 95% confluence. First, plates were washed once with ice cold 1X PBS, then cells were lysed with RIPA buffer, containing cOmplete Protease Inhibitor Cocktail, PhosStop Phosphatase Inhibitor Cocktail, and PMSF. Cell lysates were left on ice for 30 min, and vortexed every 5 min. Then, cells were stored at −80°C. Before use, samples were thawed and centrifuged at 12,000 g at 4°C for 20 min. Then, supernatant was transferred to a new Eppendorf tube. Protein concentration was measured with the DC Protein Assay, following the manufacturer’s microplate assay protocol, and using an Infinite 200 PRO microplate reader (Tecan) to measure absorbance. 25 µg protein/sample was combined with RIPA buffer, NuPAGE LDS Sample Buffer, and NuPAGE Sample Reducing Agent, and samples were incubated at 95°C for 10 min. Then, samples were run on a NuPAGE 10% Bis-Tris Gel in MOPS SDS Running Buffer. Next, proteins were transferred to an Odyssey nitrocellulose membrane. Membranes were blocked in Odyssey Blocking Buffer diluted 1:1 with PBS at room temperature for 1 hour. Membranes were incubated in primary antibody diluted in blocking buffer (1:2000 anti-GLS1; 1:1000 anti-pyruvate carboxylase) at 4°C overnight. The following day, membranes were washed with PBS-T and incubated in anti-β-actin antibody diluted in blocking buffer (1:60,000) for 1 hr at room temperature. Then, membranes were washed with PBS-T and incubated in secondary antibody diluted in blocking buffer for 45 min at room temperature (1:50,000 IRDye 680RD Donkey anti-Mouse, 1:5,000 IRDye 800CW Donkey anti-Rabbit). Membranes were washed with PBS-T again and proteins were visualized using a ChemiDoc MP Imaging System (Bio-Rad). Band intensity was quantified using Image Lab Software (Bio-Rad).

#### Targeted GC/MS

##### Sample preparation

Cells were plated in a 6-well plate at a density of 50,000 cells/well in DMEM/F-12 media supplemented with 10% FBS. 6 technical replicates were plated. 24 hrs after plating, cells were treated with CB-839 (50 nM) or vehicle (DMSO). After 24 hrs, media was removed and cells were washed on dry ice twice with ice-cold 1X PBS and scraped into 200 µL of a cold 80:20 methanol:water mixture together with 10 µL of a cocktail of labeled amino acids and TCA cycle internal standards. All internal standards concentrations were 2.5 mM, except for cystine which was 1.25 mM. Two wells were combined per one technical replicate. Next, cell suspension was centrifuged at 12,000 g for 20 min at 4°C. The supernatant was transferred to a flat bottom GC insert vial and dried at room temperature using a Labconco CentriVap Concentrator. The cell debris pellet was stored at −80°C for protein quantification. Then, the dried pellets were derivatized following the method described in (Fiehn, 2016). Briefly, the dried pellets were derivatized with 20 µL of 20 mg/mL methoxyamine HCl in pyridine and incubated on at 38°C for 90 min. Immediately following the 90 min incubation, 80 µL of MTBSTFA (with 1% t-BDMCS were added and samples were incubated at 38°C for 30 min.

To assess protein concentration in each sample, cell debris pellets were resuspended in 200 µL of RIPA lysis buffer cocktail, containing protease inhibitor cocktail, 2 mM PMSF, and 1 mM sodium orthovanadate. Cell lysates were left on ice for 30 - 60 min, and periodically vortexed vigorously. Protein concentration was measured with the Pierce BCA assay, following the manufacturer’s microplate assay protocol, and using a Synergy HT microplate reader (BioTek) to measure absorbance.

##### GC/MS analysis

Derivatized samples were run on an Agilent 7890A GC coupled to an Agilent 5975C MS and data was acquired and analyzed in MSD ChemStation E.02.02.1431 (Agilent). The GC temperature program was set to: initial temperature: 80°C, hold for 2 min; temperature rate 1: 10°C/min to 300°C, hold for 6 min. Data was collected in SIM mode (see Table S3 for ions). Metabolite peak areas in each sample were quantified relative to nmole internal standard amounts and normalized to µg protein.

#### Glutathione assay

Oxidized, reduced, and total glutathione was assessed using a spectrophotometric recycling assay. Briefly, cells were plated in 100 mm dishes at a density of 250,000 cells/dish in DMEM/F-12 media supplemented with 10% FBS. 3 technical replicates were plated. 24 hrs after plating, cells were treated with vehicle, CB-839 (50 nM), or BSO (10 µM). Cells were incubated for 24 hrs. Then, cells were scraped into 5% 5-sulfosalicylic acid and frozen. Both male and female cells were 70-80% confluent at time of cell harvest. Total glutathione was quantified using the 5,5′-dithiobis-2-nitrobenzoic acid recycling assay as described previously by Anderson (Anderson, 1985). Reduced and oxidized glutathione were distinguished using 2-vinylpyridine (2VP) mixed 1:1 (vol:vol) with ethanol as previously described by Griffith (Griffith, 1980). 20 µL of 2VP/ethanol was added for each 100 µL sample. All sample data were normalized to protein content, using the method of Lowry (Lowry et al., 1951).

#### DHE stain

Superoxide accumulation was assessed using the dihydroethidium (DHE) stain. Briefly, cells were plated in duplicates in a 96-well plate at a density of 2000 cells/well in DMEM/F-12 media supplemented with 10% FBS. 24 hrs after plating, cells were treated with vehicle, CB-839 (50 nM), or BSO (10 µM). Cells were treated for 24 hrs. Then, media was carefully removed from wells and cells were incubated in fresh culture media containing 5 µM DHE (10 mM stock solution in DMSO) at 37°C for 30 min (protected from light). Then, media was carefully removed, and cells were fixed with ice cold 4% paraformaldehyde at 4°C for 15 min. Next, plates were washed 3 times with 1X PBS and nuclei were counterstained with Hoechst 33342 (1 µg/mL in PBS). Plates were stored at 4°C protected from light until analysis. Images were acquired using an Operetta CLS High-Content Analysis System (PerkinElmer) and the Harmony High-Content Imaging and Analysis Software. DHE staining intensity in the nucleus was quantified by an individual blinded to treatment conditions by ImageJ analysis of mean fluorescent intensity. For each experiment, staining across 3 fields of view per well was averaged across 2 wells per treatment condition. Each field of view contained at least 20 cells.

#### Flow Cytometry

Cells were harvested and plated in 150 mm dishes at a density of 300,000 cells/dish in DMEM/F-12 media supplemented with 10% FBS. 24 hrs after plating, cells were treated with vehicle, CB-839 (50 nM), or BSO (10 μM). 48 hrs after treatment, cells were harvested with Accutase, counted, and aliquoted to be stained with either propidium iodide (cell cycle analysis) or Annexin V, Pacific Blue Conjugate (apoptotic cell detection). For propidium iodide staining, 1.0 x 10^6^ cells per condition were fixed for at least 15 min in 70% EtOH, washed with 1X PBS/1% FBS, strained through a 70 μm filter, and resuspended in 200 μL of propidium iodide solution containing 15 μL 500X propidium iodide, 9.9 mL 1X PBS, 100 μL 10% Triton-X-100, and 25 μL RNAse A. Cells were then incubated in the dark at 37°C for 20 min and then kept on ice (protected from light) until analysis. For Annexin V staining, cells were first resuspended at a concentration of 1.0 x 10^6^ cells in 1 mL of 1X Annexin Binding Buffer. 100 μL of each suspension was then transferred to a 5 mL flow cytometry tube, to which 5μL of Annexin V, Pacific Blue Conjugate were added. Cells were incubated for 15 min at room temperature in the dark, resuspended with an additional 400μL of 1X Annexin Binding Buffer, and kept on ice (protected from light) until analysis. All samples were run on the same ZE5 Cell Analyzer (Bio-Rad) with unstained/untreated and single positive controls. Apoptosis and cell cycle analyses were each performed three times on independent sample preparations.

FlowJo software was utilized to analyze all flow cytometry data. For cell cycle analysis, lines were drawn to demarcate which regions of each propidium iodide histogram corresponded to the following phases of the aneuploid cell cycle: G0/G1 for the diploid population, S phase for the diploid population, G2/M for the diploid population + G0/G1 for the tetraploid population (these peaks are indistinguishable), S phase for the tetraploid population, and G2/M for the tetraploid population (for representative cell cycle histograms see Fig. S4D and S5C). The percentage of the entire cell population in each phase was calculated according to the area of each demarcated region. For apoptosis analysis, Annexin V, Pacific Blue Conjugate histograms for each condition were compared by overlaying them with one another. Positive shift in staining intensity correlates with an increased number of cells stained with the Annexin V, Pacific Blue Conjugate.

### Quantification and Statistical Analysis

#### Data representation and statistical analysis

Data in Figure 1 were analyzed using R version 4.1.1 (R Core Team, 2021), the R/Bioconductor software package limma version 3.48.3 (Ritchie et al., 2015), and GraphPad Prism software version 9.0. Data were graphed using GraphPad Prism software version 9.0. For statistical analysis of sex-specific enrichment of super-pathways, sub-pathways, and individual metabolites data were analyzed using LIMMA where the moderated two sample t-test was used to compare metabolite expression between sex, and p-values were adjusted with the false discovery rate (FDR) approach using the Storey’s q-value method. For statistical analysis of nonrandom associations between male and female sample enrichment or metabolite enrichment in males or female a two-tailed Fisher’s exact test was applied.

Data in Fig. 2-7 were graphed and analyzed using GraphPad Prism software version 9.0. Data are expressed as mean ± SEM or median ± 95% confidence interval (specified in figure legends). All data shown as normalized data were normalized independently within each experiment. Each technical replicate was divided by the average of the control treatment (glutamine deprivation and drug treatments), the male value (glutamine isotope tracing experiments), or the female value (glucose isotope tracing experiments and glutamine uptake assay). For statistical analysis of two groups, test for normality distribution was first performed, using the D’Agostino-Pearson omnibus K2 test. If samples had a normal distribution an unpaired, two-tailed, parametric t-test was performed. If samples did not have a normal distribution an unpaired, two-tailed, non-parametric Mann-Whitney test was performed. Metabolite data and cell cycle analysis data were compared between two groups using two-tailed, parametric t-tests and raw p-values were adjusted using the FDR approach using the two-stage step-up method of Benjamini, Krieger, and Yekutielii. Dose-response curve concentrations were log_10_ transformed, curves were fit to normalized response versus log(inhibitor) concentration using the 4-parameters dose response logistic model, and absolute IC_50_ values were calculated. Ordinary two-way ANOVAS were performed to compare the dose response curve data between sex. For correlation analyses, nonparametric Spearman correlation coefficient was calculated, and linear regression was fit to model the linear relationship between two markers. In all cases an FDR adjusted p-value or raw p-value<0.05 was considered statistically significant.

All experiments using male and female transformed astrocytes were carried out at least three times using male and female transformed astrocytes from one cell line (one lot). All dose response-curves were carried out three times using three separately established cell lines – each consisting of a corresponding male and female line generated at the same time (three lots).

