## Supplemental Material for "Sex Differences in Brain Tumor Glutamine Metabolism Reveal Sex-Specific Vulnerabilities to Treatment"

#### Figure S1

**A**

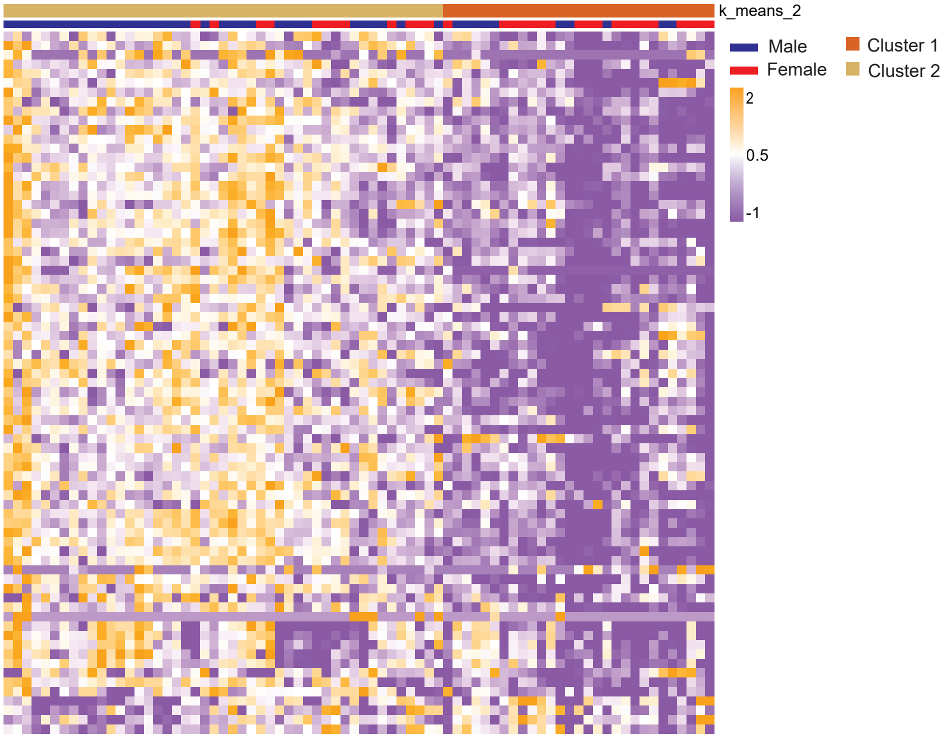


**B**

| | Cluster 1 = low metabolite abundance | Cluster 2 = high metabolite abundance |
|---|---|---|
| Male | 10 | 34 |
| Female | 19 | 13 |

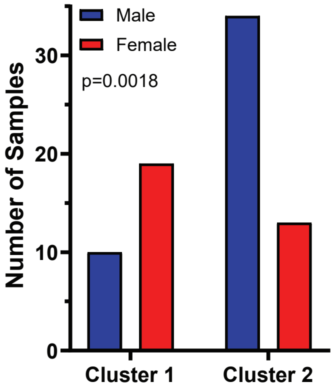

**Figure S1. Sex differences in GBM metabolite abundance parallel those found in serum, related to Figure 1. A,** K-means clustering (n=2) of the top 75 metabolites (determined by greatest mean difference) of male (n=44) and female (n=32) GBM surgical specimens. Patient specimens and metabolites are represented in columns and rows, respectively. Cluster 1 represents the low metabolite abundance group; cluster 2 represents the high metabolite abundance group. **B,** Male and female tumor sample distribution in cluster 1 and cluster 2. ***p=0.0018 (two-tailed Fisher's exact test).

#### Figure S2

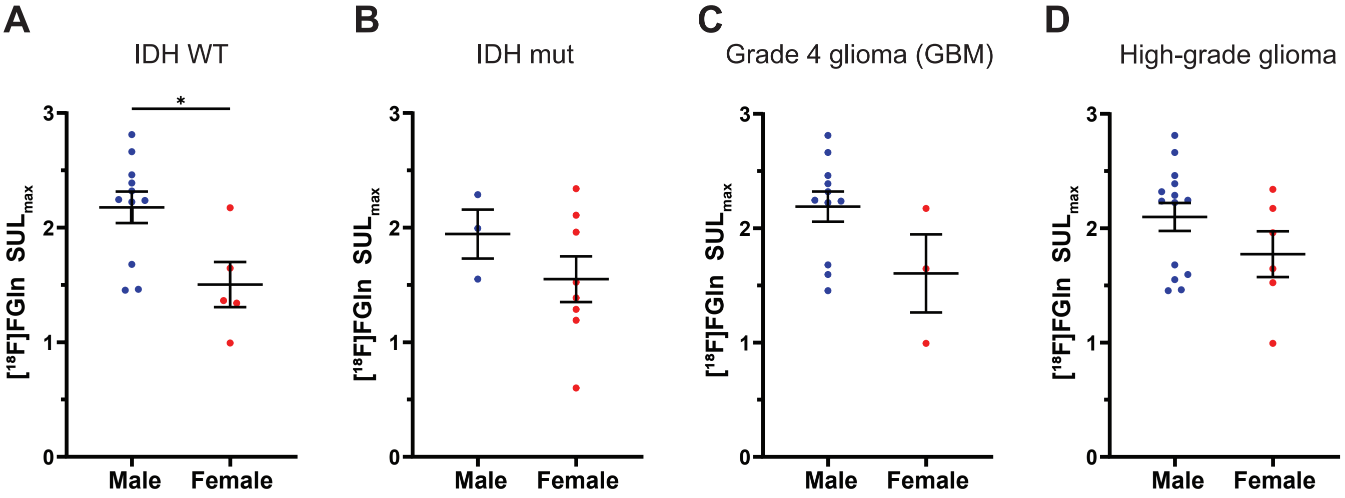
A
IDH WT
B
IDH mut
C
Grade 4 glioma (GBM)
D
High-grade glioma
[18F]FGln SULmax
*
0
1
2
3
Male
Female

**Figure S2. Glutamine uptake in male and female patient glioma subsets, related to Figure 2. A-D,** Quantification of [$^{18}$F]FGln uptake in male and female IDH wild-type (WT) gliomas (**A**),IDH mutant (mut) gliomas (**B**), grade 4 gliomas (GBM) (**C**), and high-grade gliomas (HGG) (**D**) (SULmax). Data are mean +/- SEM of n=11 male and n=5 female IDH WT patients, n=3 male and n=8 female IDH mutant patients, n=11 male and n=3 female GBM, and n=14 male and n=6 female HGG. *p<0.05 (t-test).

### Figure S3

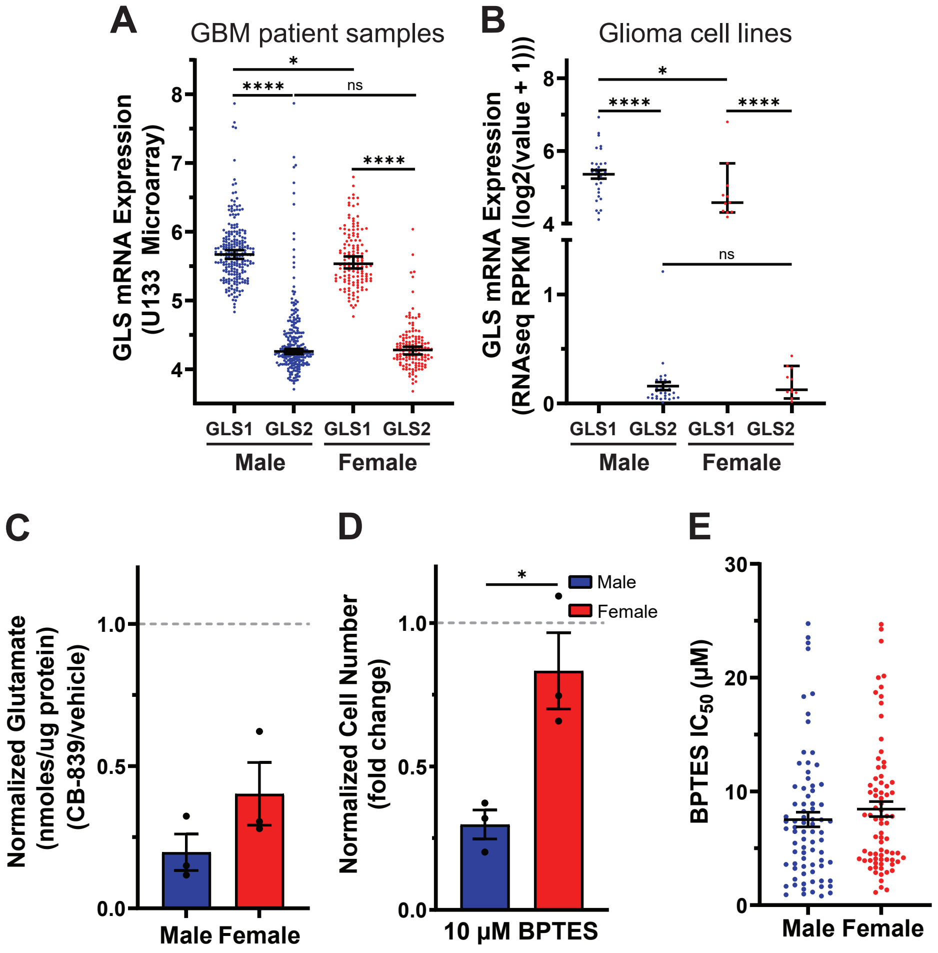
A
GBM patient samples
GLS mRNA Expression (U133 Microarray)
*
****
ns
****
GLS1 GLS2 GLS1 GLS2
Male
Female
B
Glioma cell lines
GLS mRNA Expression (RNAseq RPKM (log2(value + 1)))
*
****
****
ns
GLS1 GLS2 GLS1 GLS2
Male
Female
C
Normalized Glutamate (nmoles/ug protein) (CB-839/vehicle)
Male Female
D
Normalized Cell Number (fold change)
*
Male
Female
10 μM BPTES
E
BPTES IC50 (μM)
Male Female

**Figure S3. GLS1 expression and dependency is greater in male transformed astrocytes and gliomas, related to Figure 4. A,** GLS1 and GLS2 mRNA expression levels in n=232 male and n=139 female GBM patient samples. Data are median +/- 95% CI. ****p<0.0001 (Mann-Whitney test). **B,** GLS1 and GLS2 mRNA expression levels in n=32 male and n=11 female human glioma cell lines. Data are median +/- 95% CI. ****p<0.0001 (Mann-Whitney test). **C,** Fold change of cellular glutamate levels of transformed astrocytes treated with CB-839 for 24 hrs. Glutamate levels were measured via targeted GS/MS analysis. Data are mean of n=3/sex and treatment (three independent experiments, one male and female cell line). **D,** Cell number assay of transformed astrocytes treated with BPTES or vehicle. Data are mean +/- SEM of n=3/sex (three independent experiments, one male and female cell line). *p<0.05 (t-test). **E,** $IC_{50}$ values of male and female tumor cell lines treated with BPTES for 4 days (data obtained from Daemen et al., 2018). Data are mean +/- SEM of n = 77 male and n = 76 female tumor cell lines (t-test).

#### Figure S4

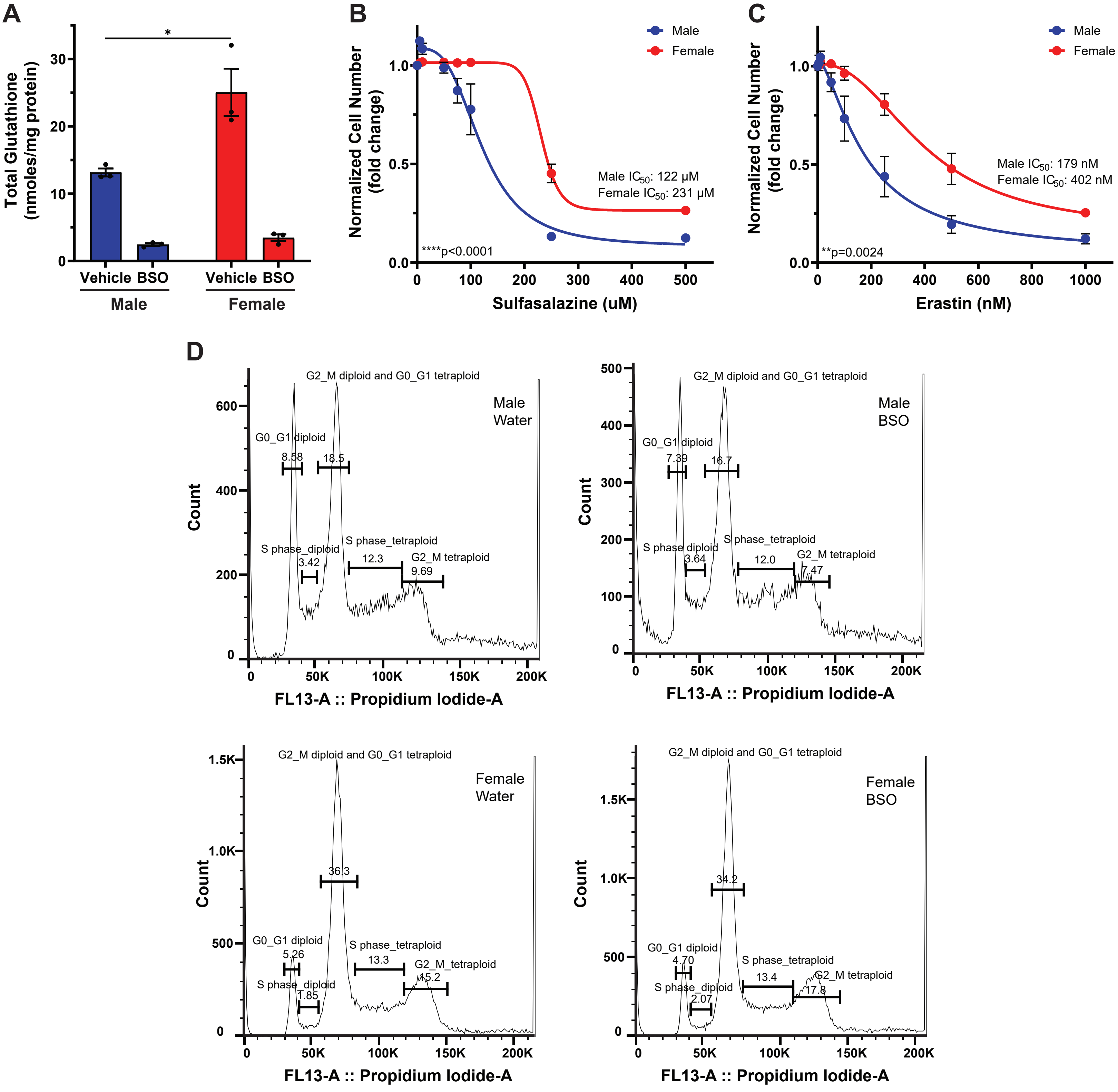
A
Total Glutathione (nmoles/mg protein)
*
Vehicle BSO
Male
Vehicle BSO
Female
B
Normalized Cell Number (fold change)
Male
Female
Male IC50: 122 μM
Female IC50: 231 μM
****p<0.0001
Sulfasalazine (uM)
C
Normalized Cell Number (fold change)
Male
Female
Male IC50: 179 nM
Female IC50: 402 nM
**p=0.0024
Erastin (nM)
D
Male Water
G2_M diploid and G0_G1 tetraploid
G0_G1 diploid
8.58
18.5
S phase_diploid
3.42
S phase_tetraploid
12.3
G2_M tetraploid
9.69
Count
FL13-A :: Propidium Iodide-A
Male BSO
G2_M diploid and G0_G1 tetraploid
G0_G1 diploid
7.39
16.7
S phase_diploid
3.64
S phase_tetraploid
12.0
G2_M tetraploid
7.47
Count
FL13-A :: Propidium Iodide-A
Female Water
G2_M diploid and G0_G1 tetraploid
36.3
G0_G1 diploid
5.26
S phase_diploid
1.85
S phase_tetraploid
13.3
G2_M_tetraploid
15.2
Count
FL13-A :: Propidium Iodide-A
Female BSO
G2_M diploid and G0_G1 tetraploid
34.2
G0_G1 diploid
4.70
S phase_diploid
2.07
S phase_tetraploid
13.4
G2_M tetraploid
17.8
Count
FL13-A :: Propidium Iodide-A

**Figure S4. Male transformed astrocytes are more dependent on glutathione for redox balance, related to Figure 5. A,** Total glutathione levels in transformed astrocytes treated with buthionine sulfoximine (BSO) or vehicle. Data are mean +/- SEM of n=3/sex (three independent experiments, one male and female cell line). *p<0.05 (t-test). **B-C,** Sulfasalazine (SAS) **(B)** and erastin **(C)** dose-response curves of transformed astrocytes. Data are mean +/- SEM of n=3/sex (three independent experiments, one male and female cell line). **p=0.0024, ****p<0.0001 (two-way ANOVA). **D**, Representative propidium iodide histograms of transformed astrocytes treated with BSO or vehicle. The cell cycle subfractions (G0/G1, S-phase, G2/M) are depicted for the diploid and tetraploid cell populations. Histograms shown are representative spectra chosen from three independent experiments.

### Figure S5

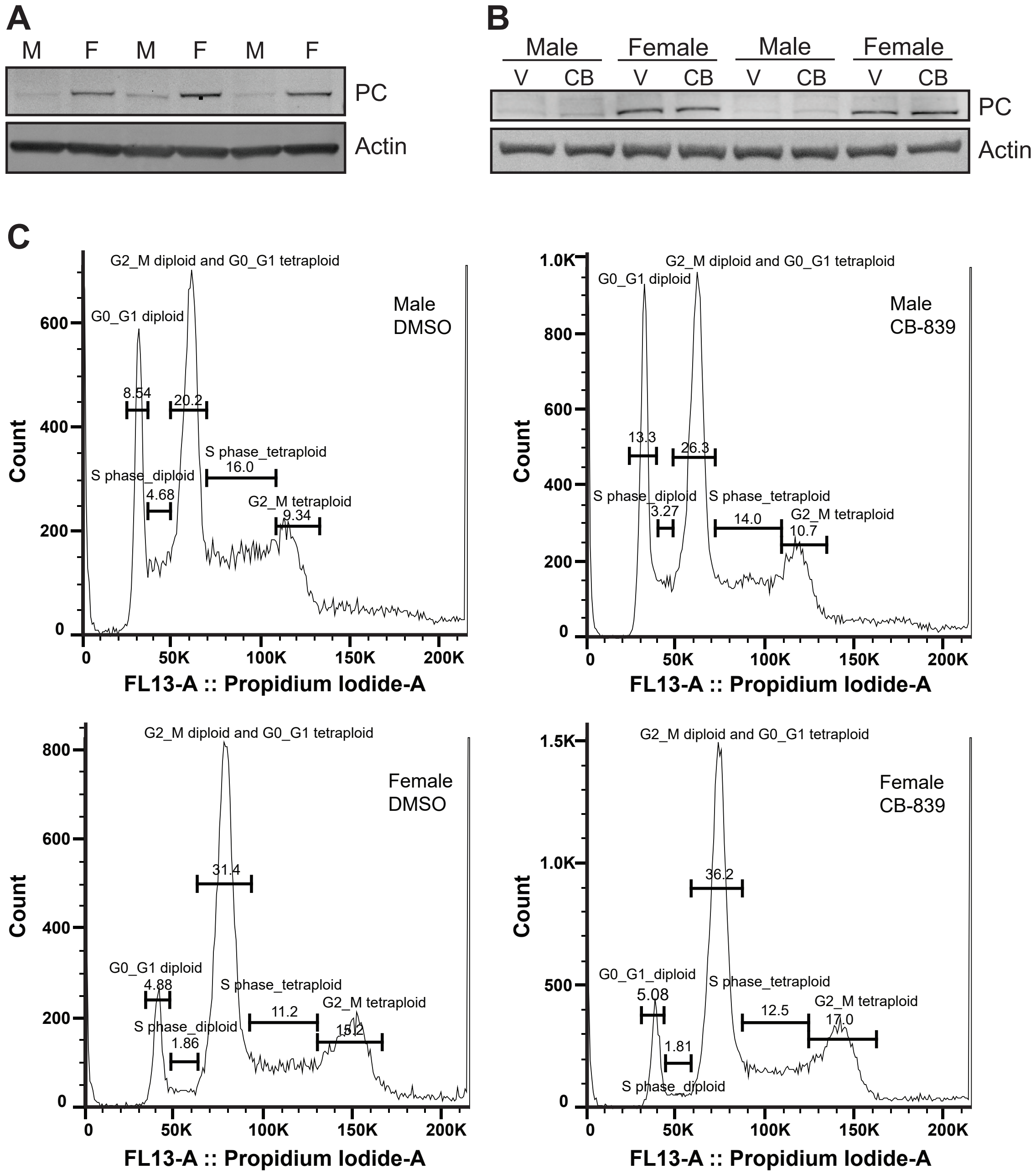
A
M
F
M
F
M
F
PC
Actin
B
Male
Female
Male
Female
V
CB
V
CB
V
CB
V
CB
PC
Actin
C
G2_M diploid and G0_G1 tetraploid
G0_G1 diploid
Male
DMSO
8.54
20.2
S phase_tetraploid
16.0
S phase_diploid
4.68
G2_M tetraploid
9.34
Count
FL13-A :: Propidium Iodide-A
0
200
400
600
50K
100K
150K
200K
G2_M diploid and G0_G1 tetraploid
G0_G1 diploid
Male
CB-839
13.3
26.3
S phase_diploid
3.27
S phase_tetraploid
14.0
G2_M tetraploid
10.7
Count
FL13-A :: Propidium Iodide-A
0
200
400
600
800
1.0K
G2_M diploid and G0_G1 tetraploid
Female
DMSO
31.4
G0_G1 diploid
4.88
S phase_tetraploid
11.2
G2_M tetraploid
15.2
S phase_diploid
1.86
Count
FL13-A :: Propidium Iodide-A
0
200
400
600
800
G2_M diploid and G0_G1 tetraploid
Female
CB-839
36.2
G0_G1_diploid
5.08
S phase_tetraploid
12.5
G2_M tetraploid
17.0
1.81
S phase_diploid
Count
FL13-A :: Propidium Iodide-A
0
500
1.0K
1.5K

**Figure S5. Male transformed astrocytes require glutamine to replenish their TCA cycle, related to Figure 6. A-B,** Western blots showing PC expression in transformed astrocytes under basal conditions (**A**) and upon treatment with CB-839 or vehicle (**B**) from three independent experiments. V=vehicle, CB=CB-839. **C**, Representative propidium iodide histograms of transformed astrocytes treated with CB-839 or vehicle. The cell cycle subfractions (G0/G1, S-phase, G2/M) are depicted for the diploid and tetraploid cell population. Histograms shown are representative spectra chosen from three independent experiments.

**Table S1. Glioma patient characteristics, related to Figure 2**

| Characteristic | Female, N = 13[a] | Male, N = 15[a] | Total, N = 28[a] |
|---|---|---|---|
| **Diagnosis** | | | |
| Anaplastic astrocytoma | 3 (23%) | 3 (20%) | 6 (21%) |
| Anaplastic ependymoma | 1 (8%) | 0 (0%) | 1 (4%) |
| Anaplastic oligodendroglioma | 2 (15%) | 0 (0%) | 2 (7%) |
| Astrocytoma | 4 (31%) | 0 (0%) | 4 (14%) |
| Glioblastoma | 3 (23%) | 11 (73%) | 14 (50%) |
| Oligodendroglioma | 0 (0%) | 1 (7%) | 1 (4%) |
| **Grade** | | | |
| 2 | 4 (31%) | 1 (7%) | 5 (18%) |
| 3 | 6 (46%) | 3 (20%) | 9 (32%) |
| 4 | 3 (23%) | 11 (73%) | 14 (50%) |
| **IDH status** | | | |
| Mutant | 8 (62%) | 3 (20%) | 11 (39%) |
| WT | 5 (38%) | 11 (73%) | 16 (57%) |
| Unknown | 0 (0%) | 1 (7%) | 1 4%) |

[a]Statistics presented: n (%)

**Table S2. m/z values monitored for glutamine and glucose labeling studies, related to Figure 3 and Figure 6**

| Metabolite | m/z monitored |
|---|---|
| **GC/MS [$^{13}C_5{}^{15}N_2$]Gln** | |
| Alanine | 260.1, 261.1, 262.1, 263.1, 264.1, 265.1, 266.1, 267.1 |
| Fumarate | 287.1, 288.1, 289.1, 290.1, 291.1, 292.1, 293.1, 294.1 |
| Valine | 288.1, 289.1, 290.1. 291.1, 292.1, 293.1, 294.1, 295.1, 296.1, 297.1 |
| Succinate | 289.1, 290.1, 292.1, 293.1, 294.1, 295.1, 296.1 |
| Leucine | 302.1, 303.1, 304.1, 305.1, 306.1, 307.1, 308.1, 309.1, 310.1, 311.1, 312.1 |
| Isoleucine | 302.1, 303.1, 304.1, 305.1, 306.1, 307.1, 308.1, 309.1, 310.1, 311.1, 312.1 |
| α-Ketoglutarate | 346.2, 347.2, 348.2, 349.2, 350.2, 351.2, 352.2, 353.2, 354.2 |
| Serine | 390.2, 391.2, 392.2, 393.2, 394.2, 395.2, 396.2, 397.2 |
| Aspartate | 418.3, 419.3, 420.3, 421.3, 422.3, 423.3, 424.3, 425.3, 426.3 |
| Malate | 419.3, 420.3, 421.3, 422.3, 423.3, 424.3, 425.3. 426.3 |
| Glutamate | 432.3, 433.3, 434.3, 435.3, 436.3, 437.3, 438.3, 439.3, 440.3, 441.3 |
| Citrate | 591.4, 592.4, 593.4, 594.4, 595.4, 596.4, 597.4, 598.4, 599.4, 600.4 |
| **LC/MS [$^{13}C_5{}^{15}N_2$]Gln** | |
| Glutathione (red.) | 306.1, 307.1, 308.1, 309.1, 310.1, 311.1, 312.1, 313.1, 314.1, 315.1, 316.1 |
| Glutathione (oxid.) | 611.1, 612.1, 613.2, 614.2, 615.2, 616.2, 617.2, 618.2, 619.2, 620.2, 621.2, 622.2, 623.2, 624.2, 625.2, 626.2, 627.2, 628.2, 629.2, 630.2, 631.2 |
| **GC/MS [$^{13}C_6$]Glc** | |
| Fumarate | 287.1, 288.1, 289.1, 290.1, 291.1, 292.1, 293.1 |
| Aspartate | 418.3, 419.3, 420.3, 421.3, 422.3, 423.3, 424.3 |
| Malate | 419.3, 420.3, 421.3, 422.3, 423.3, 424.3, 425.3 |
| Citrate | 591.4, 592.4, 593.4, 594.4, 595.4, 596.4, 597.4 |

**Table S3. Metabolite ions monitored for targeted GC/MS analysis, related to Figure 5**

| Metabolites | Ions | Internal Standards (IS) | IS Ions |
|---|---|---|---|
| Alanine | 232.1 | $^{13}C_3{}^{15}N$ L-Alanine | 235.1 |
| Glycine | 218.1 | $^{13}C_2{}^{15}N$ Glycine | 220.1 |
| Valine | 186.2 | $^{13}C_5{}^{15}N$ L-Valine | 191.2 |
| Leucine | 200.2 | $^{13}C_6{}^{15}N$ L-Leucine | 206.2 |
| Isoleucine | 200.2 | $^{13}C_6{}^{15}N$ L-Isoleucine | 206.2 |
| Proline | 184.1 | $^{13}C_5{}^{15}N$ L-Proline | 189.1 |
| Methionine | 218.1 | $^{13}C_5{}^{15}N$ L-Methionine | 223.2 |
| Serine | 288.1 | $^{13}C_3{}^{15}N$ L-Serine | 291.1 |
| Threonine | 303.2 | $^{13}C_4{}^{15}N$ L-Threonine | 306.2 |
| Phenylalanine | 302.1 | $^{13}C_9{}^{15}N$ L-Phenylalanine | 305.1 |
| Aspartate | 302.1 | $^{13}C_4{}^{15}N$ L-Aspartate | 305.1 |
| Cysteine | 302.1 | $^{13}C_3{}^{15}N$ L-Cysteine | 305.1 |
| Glutamate | 432.3 | $^{13}C_5{}^{15}N$ L-Glutamate | 438.3 |
| Asparagine | 417.3 | $^{13}C_4{}^{15}N_2$ L-Asparagine | 423.3 |
| Lysine | 300.2 | $^{13}C_6{}^{15}N_2$ L-Lysine | 307.2 |
| Glutamine | 431.3 | $^{13}C_5{}^{15}N_2$ L-Glutamine | 438.3 |
| Histidine | 196.1 | $^{13}C_6{}^{15}N_3$ L-Histidine | 202.2 |
| Tyrosine | 302.1 | $^{13}C_9{}^{15}N$ L-Tyrosine | 305.1 |
| Tryptophan | 302.1 | $^{13}C_{11}{}^{15}N_2$ L-Tryptophan | 305.1 |
| Cystine | 348.2 | $^{13}C_6{}^{15}N_2$ L-Cystine | 352.2 |
| Pyruvate | 174.0 | $^{13}C_3$ Pyruvate | 177.0 |
| Lactate | 261.1 | $^{13}C_3$ Lactate | 264.1 |
| Citrate | 459.3 | $^{13}C_6$ Citrate | 465.3 |
| α-ketoglutarate | 346.1 | $^{13}C_4$ α-ketoglutarate | 350.1 |
| Succinic acid | 289.1 | $^{13}C_4$ Succinic acid | 293.1 |
| Fumaric acid | 287.1 | $^{13}C_4$ Fumaric acid | 291.1 |
| Malic acid | 419.3 | $^{13}C_4$ Malic acid | 423.3 |

**KEY RESOURCES TABLE**

| REAGENT or RESOURCE | SOURCE | IDENTIFIER |
|---|---|---|
| Antibodies | | |
| anti-GLS1 (KGA/GAC) polyclonal antibody | Proteintech | Cat# 12855-1-AP; RRID:AB_2110381 |
| anti-pyruvate carboxylase polyclonal antibody | Proteintech | Cat# 16588-1-AP; RRID:AB_1851513 |
| anti-β-actin monoclonal antibody | Sigma | Cat# A1978; RRID:AB_476692 |
| IRDye 680RD donkey anti-mouse | LI-COR | Cat# 926-68072; RRID:AB_10953628 |
| IRDye 800CW donkey anti-rabbit | LI-COR | Cat# 926-32213; RRID:AB_621848 |
| Chemicals, peptides, and recombinant proteins | | |
| [$^{18}$F]fluoroglutamine | Memorial Sloan Kettering Cancer Center's Radiochemistry and Molecular Imaging Probe Core Facility | N/A |
| CB-839 | Selleckchem | Cat# S7655 |
| BPTES | Selleckchem | Cat# S7753 |
| Menadione | Selleckchem | Cat# S1949 |
| Buthionine sulfoximine | Sigma | Cat# 5.08228 |
| DMEM/F-12 | USBiological | Cat# D9807-06 |
| Dialyzed FBS | Gibco | Cat# 26400044 |
| D-glucose | Corning | Cat# 25-037-CI |
| L-glutamine | Gibco | Cat# 25030081 |
| Sulforhodamine B | Sigma | Cat# S1402 |
| $H_2O_2$ | Sigma | Cat# H1009 |
| N-acetyl-cysteine | Sigma | Cat# A9165 |
| Dimethyl-α-ketoglutarate | Sigma | Cat# 349631 |
| Sodium pyruvate | Sigma | Cat# P5280 |
| [$^{13}C_6$]glucose | Cambridge Isotope Laboratories | Cat# CLM-1396-PK |
| [$^{13}C_5{}^{15}N_2$]glutamine | Cambridge Isotope Laboratories | Cat# CNLM-1275-H-PK |
| Methanol (MS grade) | Thermo Fisher Scientific | Cat# A456 |
| Water (MS grade) | Sigma | Cat# 900682 |
| Acetonitrile (MS grade) | Sigma | Cat# 900667 |
| Methoxyamine hydrochloride | Sigma | Cat# 226904 |
| Pyridine anhydrous 99.8% | Sigma | Cat# 270970 |
| MTBSTFA (with 1% t-BDMCS) | Sigma | Cat# M-108 |
| cOmplete protease inhibitor cocktail | Roche | Cat# 11697498001 |
| PhosStop phosphatase inhibitor cocktail | Roche | Cat# 4906845001 |
| NuPAGE LDS sample buffer | Thermo Fisher | Cat# NP0007 |
| NuPAGE sample reducing agent | Thermo Fisher | Cat# NP0004 |
| MOPS SDS running buffer | Thermo Fisher | Cat# NP0001 |
| Odyssey nitrocellulose membrane | LI-COR | Cat# 926-31092 |

| | | |
|---|---|---|
| Intercept blocking buffer PBS | LI-COR | Cat# 927-70003 |
| Hygromycin B | Goldbio | Cat# H-270-1 |
| D-luciferin firefly | Biosynth Carbosynth | Cat# L-8220 |
| hEGF | Sigma | Cat# E9644 |
| Matrigel basement membrane matrix | Corning | Cat# 354234 |
| [$^{13}C_6$]citric acid | Cambridge Isotope Laboratories | Cat# CLM-9021-PK |
| [$^{13}C_4$]α-ketoglutaric acid sodium salt | Cambridge Isotope Laboratories | Cat# CLM-4442 |
| [$^{13}C_4$]succinic acid | Cambridge Isotope Laboratories | Cat# CLM-1571 |
| [$^{13}C_4$]fumaric acid | Cambridge Isotope Laboratories | Cat# CLM-1529 |
| [$^{13}C_4$]L-malic acid | Cambridge Isotope Laboratories | Cat# CLM-8065 |
| Stable isotope labeled canonical amino acid mix | Cambridge Isotope Laboratories | Cat# MSK-CAA-1 |
| RIPA lysis buffer cocktail | Santa Cruz | Cat# sc-24948 |
| 5-sulfosalicylic acid | Sigma | Cat# S7408 |
| Dihydroethidium stain | Thermo Fisher | Cat# D11347 |
| Hoechst 33342 | Thermo Fisher Scientific | Cat# 62249 |
| Propidium iodide | Sigma | Cat# P4170 |
| Annexin V, pacific blue conjugate | Thermo Fisher Scientific | Cat# A35122 |
| Triton-X-100 | Sigma | Cat# 9002-93-1 |
| RNAse A, DNase and protease-free | Thermo Fisher Scientific | Cat# EN0531 |
| Annexin binding buffer | Thermo Fisher Scientific | Cat# V13246 |
| Critical commercial assays | | |
| DC protein assay kit | Bio-Rad | Cat# 5000111 |
| Pierce BCA assay kit | Thermo Fisher Scientific | Cat# 23225 and Cat# 23227 |
| Experimental models: Cell lines | | |
| Nf1-/-; DNp53 astrocytes | Rubin Laboratory | N/A |
| Software and algorithms | | |
| R v4.1.1 | R Core Team, 2021 | https://www.r-project.org/ |
| Limma v3.48.3 | Bioconductor | https://bioconductor.org/packages/release/bioc/html/limma.html DOI:10.18129/B9.bioc.limma |
| PET VCAR software | GE Healthcare | https://www.gehealthcare.com/products/advanced-visualization/all-applications/pet-vcar |

| | | |
|---|---|---|
| GraphPad Prism v9 | GraphPad | https://www.graphpad.com/scientific-software/prism/; RRID:SCR_002798 |
| Skyline software | MacLean et al., 2009 | https://skyline.ms/project/home/software/Skyline/begin.view |
| ChemStation E.02.02.1431 | Agilent | https://www.agilent.com/en/product/software-informatics/analytical-software-suite |
| FluxFix: Isotopologue Analysis Tool v0.1 | Trefely et al., 2016 | http://fluxfix.science/ |
| Image Lab Software v6.1 | Bio-Rad | https://www.bio-rad.com/en-us/product/image-lab-software?ID=KRE6P5E8Z; RRID:SCR_014210 |
| Morpheus | Broad Institute | https://software.broadinstitute.org/morpheus |
| Harmony High-Content Imaging and Analysis Software v4.5 | PerkinElmer | https://www.perkinelmer.com/product/harmony-4-9-office-license-hh17000010 |
| ImageJ v1.53a | Schneider et al., 2012 | https://imagej.nih.gov/ij/index.html; RRID:SCR_003070 |
| FlowJo v10.7.2 | FlowJo | https://www.flowjo.com/solutions/flowjo/downloads; RRID:SCR_008520 |
| Other | | |
